# MeCP2 binds to methylated DNA independently of phase separation and heterochromatin organisation

**DOI:** 10.1101/2023.05.09.539985

**Authors:** Raphaël Pantier, Megan Brown, Sicheng Han, Katie Paton, Stephen Meek, Thomas Montavon, Toni McHugh, David A. Kelly, Tino Hochepied, Claude Libert, Thomas Jenuwein, Tom Burdon, Adrian Bird

## Abstract

Correlative evidence has suggested that DNA methylation promotes the formation of transcriptionally silent heterochromatin. Accordingly, the methyl-CpG binding domain protein MeCP2 is often portrayed as a constituent of heterochromatin. This interpretation has been reinforced by the use of mouse cells as an experimental system for studying the mammalian epigenome, as heterochromatin, DNA methylation and MeCP2 colocalise in prominent foci. The findings presented here revise this view. We show that focal localisation of MeCP2 in mice is independent of heterochromatin, as DNA methylation-dependent MeCP2 foci persist even when the signature heterochromatin histone mark H3K9me3 is absent and heterochromatin protein HP1 is diffuse. Contrary to the proposal that MeCP2 forms condensates at mouse heterochromatic foci via liquid-liquid phase transition, the short methyl-CpG binding domain, which lacks the disordered domains thought to be required for condensation, is sufficient to target foci in mouse cells. Importantly, we find that the formation of MeCP2 foci in mice is highly atypical, as they are indetectable in 14 out of 16 other mammalian species, including humans. Notably, MeCP2 foci are absent in *Mus spretus* which can interbreed with *Mus musculus* but lacks its highly methylated pericentric satellite DNA repeats. We conclude that MeCP2 has no intrinsic tendency to form nuclear condensates and its localisation is independent of heterochromatin formation. Instead, the distribution of MeCP2 in the nucleus is primarily determined by global DNA methylation patterns and is typically euchromatic.

## Introduction

MeCP2 is an epigenetic regulator which controls gene expression by recognising both methylated ‘CG’ and ‘CA’ sequences (Lewis *et al*, 1992; Guo *et al*, 2014; Chen *et al*, 2015; Gabel *et al*, 2015; Kinde *et al*, 2016; Lagger *et al*, 2017; Cholewa-Waclaw *et al*, 2019; Boxer *et al*, 2020; Tillotson *et al*, 2021). MeCP2 function is particularly important in neurons (Guy *et al*, 2001; Chen *et al*, 2001; Shahbazian *et al*, 2002; Skene *et al*, 2010; Ross *et al*, 2016) and its mutation in humans is responsible for Rett syndrome, a severe neurological disorder primarily affecting females (Amir *et al*, 1999). Experiments in cell lines and transgenic mice indicate that MeCP2 acts as a molecular “bridge” between methylated DNA and the co-repressor complex NCoR to restrain expression of large numbers of neuronal genes (Lyst *et al*, 2013; Tillotson *et al*, 2017; Koerner *et al*, 2018). This model is supported by the observation that missense mutations in the methyl-DNA binding domain (MBD) and NCoR-interaction domain (NID) of MeCP2 give rise to Rett syndrome (Lyst & Bird, 2015) and by evidence that a mini-MeCP2 comprising only the MBD and NID rescues Rett-like neurological phenotypes in mice (Tillotson *et al*, 2017).

In mouse cells, MeCP2 is enriched within large foci of pericentric heterochromatin (Lewis *et al*, 1992), also known as chromocenters (Brändle *et al*, 2022), which contain heavily methylated satellite DNA repeats (Thakur *et al*, 2021). Recently, several studies have proposed that colocalization with heterochromatic foci is caused by an intrinsic tendency of MeCP2 to form “condensates” via a process of liquid-liquid phase separation (Li *et al*, 2020; Wang *et al*, 2020; Fan *et al*, 2020; Jiang *et al*, 2021; Zhang *et al*, 2022). Distinguishing phase separation from alternative origins of intracellular “membraneless compartments” *in vivo* is challenging (McSwiggen *et al*, 2019; Musacchio, 2022; Mittag & Pappu, 2022). In particular, whether MeCP2 forms phase separated condensates or simply localises to regions enriched in DNA methylation is an open question. In this study, we aimed to determine the molecular determinants of MeCP2 foci in mouse cells using MeCP2 mutant constructs and genetically engineered cell lines lacking DNA methylation or organised heterochromatin. Furthermore, we investigated the influence of genome architecture on MeCP2 sub-nuclear distribution by performing live-cell imaging across 16 different mammalian species. Our results challenge the hypothesis that MeCP2 accumulation or heterochromatin formation require phase separation in live cells and suggest that the prominent chromocenters observed in mouse cells are governed by atypical genomic features which are not present in most mammalian species.

## Results

### Only the methyl-CpG binding domain of MeCP2 is required for localisation to DNA-dense foci in mouse cells

The two best-characterised macromolecular interaction partners of MeCP2 are methylated DNA, which binds to the methyl-CpG binding domain (MBD: (Nan *et al*, 1993; Ho *et al*, 2008) and the NCoR co-repressor complex which binds to NCoR-interaction domain (NID; (Lyst *et al*, 2013); see Figure 1A). Previous work demonstrated that the MBD is essential for localisation to heavily methylated DNA-dense foci in mouse cells (Nan *et al*, 1996; Kudo *et al*, 2003; Kumar *et al*, 2008) and we confirmed this by transfection of mouse fibroblasts (NIH 3T3 cells) with constructs expressing EGFP-tagged mouse MeCP2 (Figure 1B, 1C). In these and subsequent experiments we monitored the distribution of exogenous MeCP2 in living cells using super-resolution microscopy. Deletion of 27 amino acids within the MBD (Δ99-125) (Nan *et al*, 1996) greatly decreased co-localisation with DNA-dense foci, causing instead accumulation of MeCP2 in larger non-overlapping nuclear bodies (Figure 1B, 1C). To test whether the MBD was sufficient for correct subnuclear localisation we expressed an EGFP-tagged 85 amino acid peptide corresponding to the minimal MBD (Nan *et al*, 1993).

**Figure 1.**
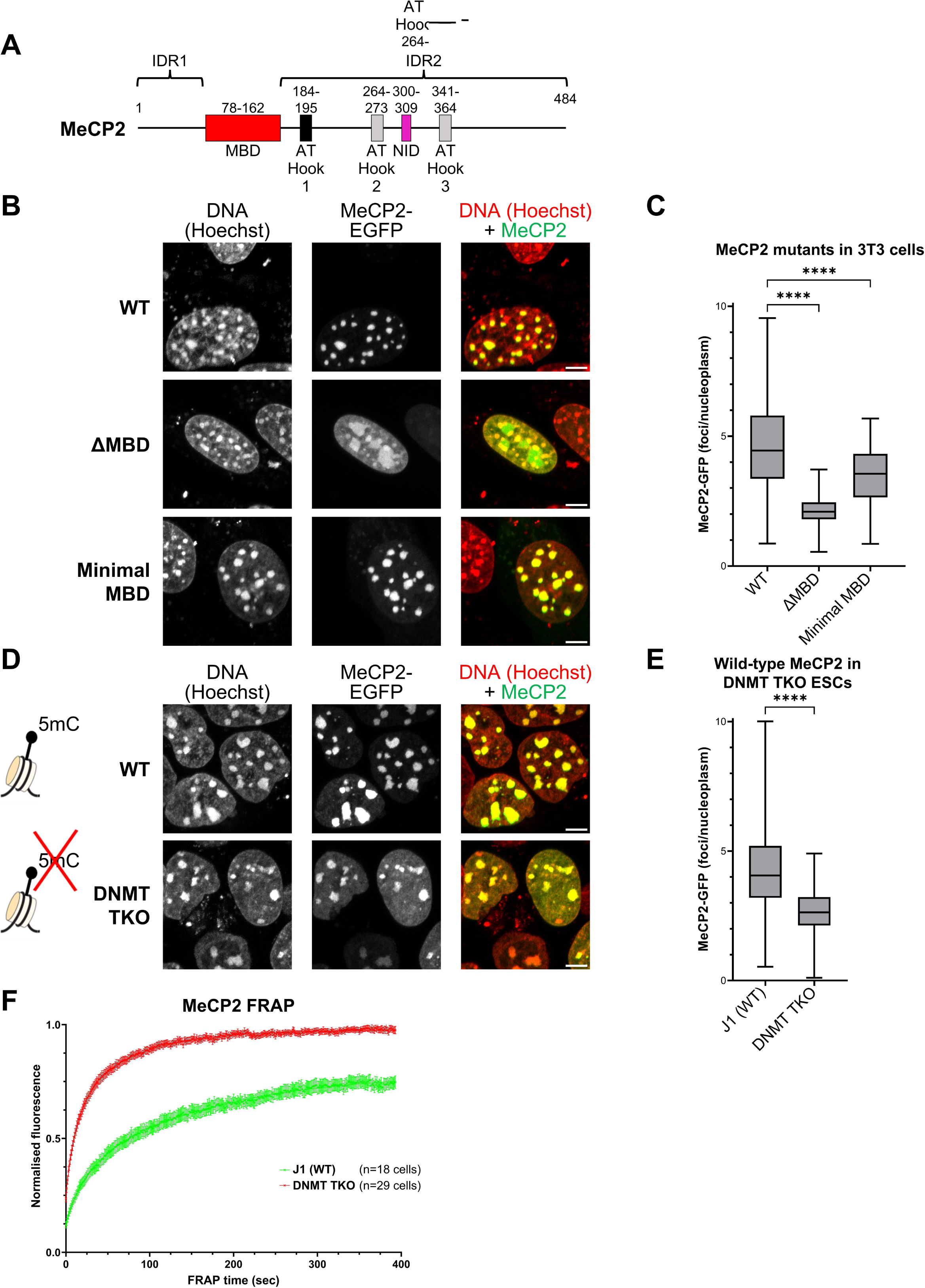
MeCP2 localises to DNA-dense foci in mouse cells via its Methyl-DNA Binding Domain (MBD) **A.** Diagram showing the structure of wild-type MeCP2 with annotated domains. MBD: Methyl-CpG Binding Domain; NID: NCoR-Interaction Domain; IDR: Intrinsically Disordered Region. **B.** Live-cell imaging of 3T3 cells transfected with the EGFP-MeCP2 constructs (see Figure S1). Hoechst staining was used to visualise DNA. Scale bars: 5µm. **C.** Box plot showing the quantification of MeCP2 wild-type and mutant fluorescence at DNA-dense foci (relative to nucleoplasm) in 3T3 cells, as described in panel B. **D.** Live-cell imaging of wild-type (J1) and *DNMT TKO* ESCs transfected with wild-type EGFP-MeCP2. Hoechst staining was used to visualise DNA. Scale bars: 5µm. **E.** Box plot showing the quantification of MeCP2 fluorescence at DNA-dense foci (relative to nucleoplasm) in wild-type (J1) and *DNMT TKO* ESCs, as described in panel A. **F.** Graph showing the FRAP quantification of wild-type EGFP-MeCP2 in wild-type (J1, green) and *DNMT TKO* (red) ESCs. Error bars: SEM.

Despite lacking the two intrinsically disordered regions of MeCP2 (IDR1 and IDR2), the minimal MBD localised to DNA-dense foci (Figure 1B). To quantify this distribution, we compared the intensity of fluorescence within foci versus the remaining nucleoplasm. This confirmed the preference of the MBD for foci, although at a slightly lower level than that of full length MeCP2 (Figure 1C). This modest reduction in binding efficiency may be due to absence of the MeCP2 nuclear localisation signal, which is not essential for MeCP2 entry into the nucleus, but may improve efficiency (Lyst *et al*, 2018). Further DNA binding specificity is may be provided by three potential AT Hooks (Klose *et al*, 2005; Baker *et al*, 2013), of which only AT Hook1 shows a marked preference for AT-rich DNA *in vitro* (Lyst *et al*, 2016) (Figure 1A). However, mutation of AT Hook1 (R188G, R190G) in a wild-type or mutant MBD molecule revealed no detectable contribution to MeCP2 sub-nuclear localisation (Figure S1A, S1B, S1C). These findings were not confined to fibroblasts, as similar results were obtained in mouse embryonic stem cells (ESCs) (Figure S2A, S2B). Taken together, the evidence suggests that the MBD is both necessary and sufficient for robust localisation to mouse heterochromatin, whereas AT-hook1 and “intrinsically disordered regions” are neither necessary nor sufficient.

### Intact heterochromatin is not required for stable MeCP2 binding to nuclear foci in mouse cells

To determine the epigenomic requirements for MeCP2 distribution inside the nucleus, we first used embryonic stem cells lacking all three DNA methyltransferases: DNMT1, DNMT3A and DNMT3B (Tsumura *et al*, 2006). These triple knockout (*DNMT TKO*) cells contain no detectable 5-methylcytosine, as confirmed by mass spectrometry of genomic DNA (Figure S3A). In agreement with a previous report (Fan *et al*, 2020), MeCP2 retained localisation to chromocenters in live *DNMT TKO* ESCs, although nucleoplasmic signal was also significantly elevated compared to the parental cell line (Figure 1D, 1E). In fact, MeCP2 and Hoechst displayed closely similar patterns, implying generalised DNA binding in the absence of DNA methylation. Neither the MBD alone nor a full-length MeCP2 lacking a functional MBD localised to chromocenters in the absence of DNA methylation (Figure S3B, S3C) .

Thus, both the MBD and non-MBD regions of MeCP2 are needed for heterochromatic localisation in the absence of DNA methylation, but neither is sufficient. These findings raised the possibility that MeCP2 localisation in the absence of DNA methylation is relatively non-specific, in which case chromatin binding dynamics would be expected to dramatically increase. To test this prediction, we performed fluorescence recovery after photobleaching (FRAP) of wild-type MeCP2 at heterochromatic foci. In line with previous studies (Klose *et al*, 2005; Kumar *et al*, 2008; Agarwal *et al*, 2011; Tillotson *et al*, 2021), wild-type ESCs showed incomplete fluorescence recovery, even >6 minutes after bleaching, whereas MeCP2 recovery was complete and rapid in *DNMT TKO* ESCs (Figure 1F, S3D). Incomplete recovery and failure of fluorescence to plateau in wild-type cells prevents simple numerical comparison between wild-type and mutant cells, but it is evident that the time to reach 50% recovery is greatly reduced in *DNMT TKO* cells (∼10 seconds versus ∼75 seconds; Table 1). The data show that the stably bound fraction of MeCP2 is abolished in the absence of DNA methylation and MeCP2 binding becomes much more transient and dynamic, suggesting a loss of DNA binding specificity.

**Table 1. MeCP2 FRAP analysis in mutant cell lines**

**Table 1.**
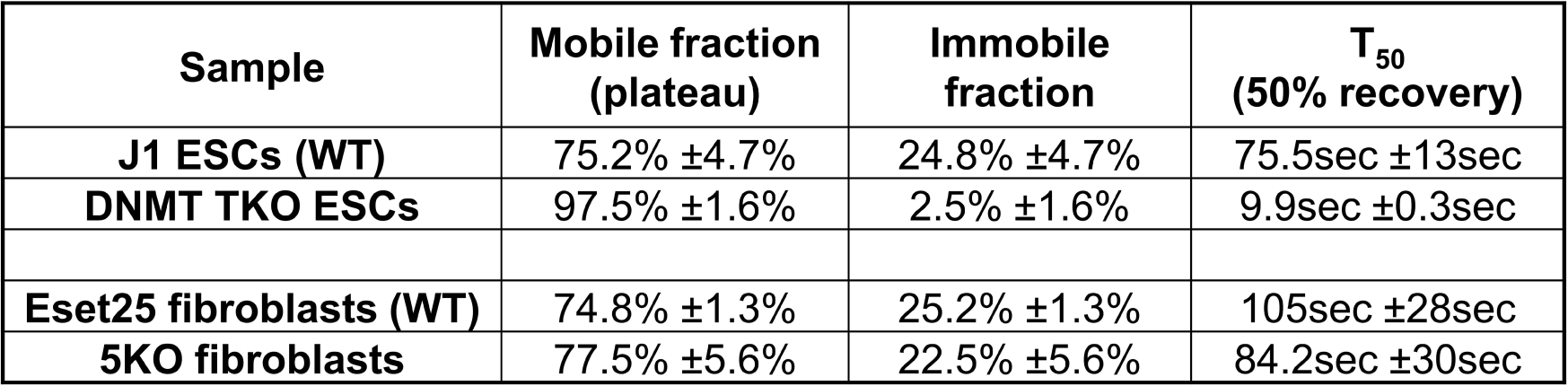
(related to Figure 1 & 2) -MeCP2 FRAP analysis in mutant cell lines

| **Sample** | **Mobile fraction**<br>**(plateau)** | **Immobile**<br>**fraction** | **T<sub>50</sub>**<br>**(50% recovery)** |
| --- | --- | --- | --- |
| **J1 ESCs (WT)** | 75.2% ±4.7% | 24.8% ±4.7% | 75.5sec ±13sec |
| **DNMT TKO ESCs** | 97.5% ±1.6% | 2.5% ±1.6% | 9.9sec ±0.3sec |
| **Eset25 fibroblasts (WT)** | 74.8% ±1.3% | 25.2% ±1.3% | 105sec ±28sec |
| **5KO fibroblasts** | 77.5% ±5.6% | 22.5% ±5.6% | 84.2sec ±30sec |

The DNA-dense foci in mouse cells at which MeCP2 accumulates correspond to pericentric heterochromatin, which is marked by trimethylation of histone H3, lysine 9 (H3K9me3). To evaluate whether MeCP2 co-localisation depends on intact heterochromatin organisation, we used mouse cell lines in which histone lysine methyltransferase genes have been deleted (Montavon *et al*, 2021). *Suv39h1/2* double knock-out (*2KO*) fibroblasts lack H3K9 trimethylation, whereas in *5KO* fibroblasts, five out of six known H3K9 methyltransferases were deleted (Suv39h1/2, Eset2, and G9a/Glp) as *Eset1* knockout is lethal (Montavon *et al*, 2021). Of note, DNA methylation is largely unaffected in these cells. Immunostaining of fixed cells confirmed loss of H3K9me3 and diffuse localisation of heterochromatin protein 1α (HP1α), both in fixed *2KO* and *5KO* fibroblasts (Figure S4A, S4B). To visualise heterochromatin in live cells, we co-transfected our cell lines with a reporter that expressed a dimerised HP1β (CBX1) chromodomain fused to a fluorescent protein (CHD-mCherry) (Figure 2A) (Villaseñor *et al*, 2020). As expected, CHD-mCherry signal became diffuse in *2KO* and *5KO* cell lines (Figure 2B). Strikingly, the nuclear distribution of full-length MeCP2 (Figure 2B, 2C) and of the MBD alone (Figure S4C, S4D) appeared unaltered by the dissolution of heterochromatin in these cells. Furthermore, FRAP analysis showed no difference in fluorescence recovery between the *5KO* and wild-type cell lines (Figure 2D, S4E, S4F and Table 1), suggesting identical dynamics of MeCP2 binding whether heterochromatin is intact or impaired. To test whether loss of MeCP2 has a direct impact on pericentric heterochromatin organisation, we performed immunofluorescence analysis in *Mecp2* knockout fibroblasts (Guy *et al*, 2001) (Figure S5A). H3K9me3-positive (Figure S5B) and HP1α-positive (Figure S5C) foci co-localising with DNA were indistinguishable from the wild-type control. The results suggest that, despite their co-location, MeCP2 accumulation and heterochromatin organisation are mutually independent phenomena.

**Figure 2.**
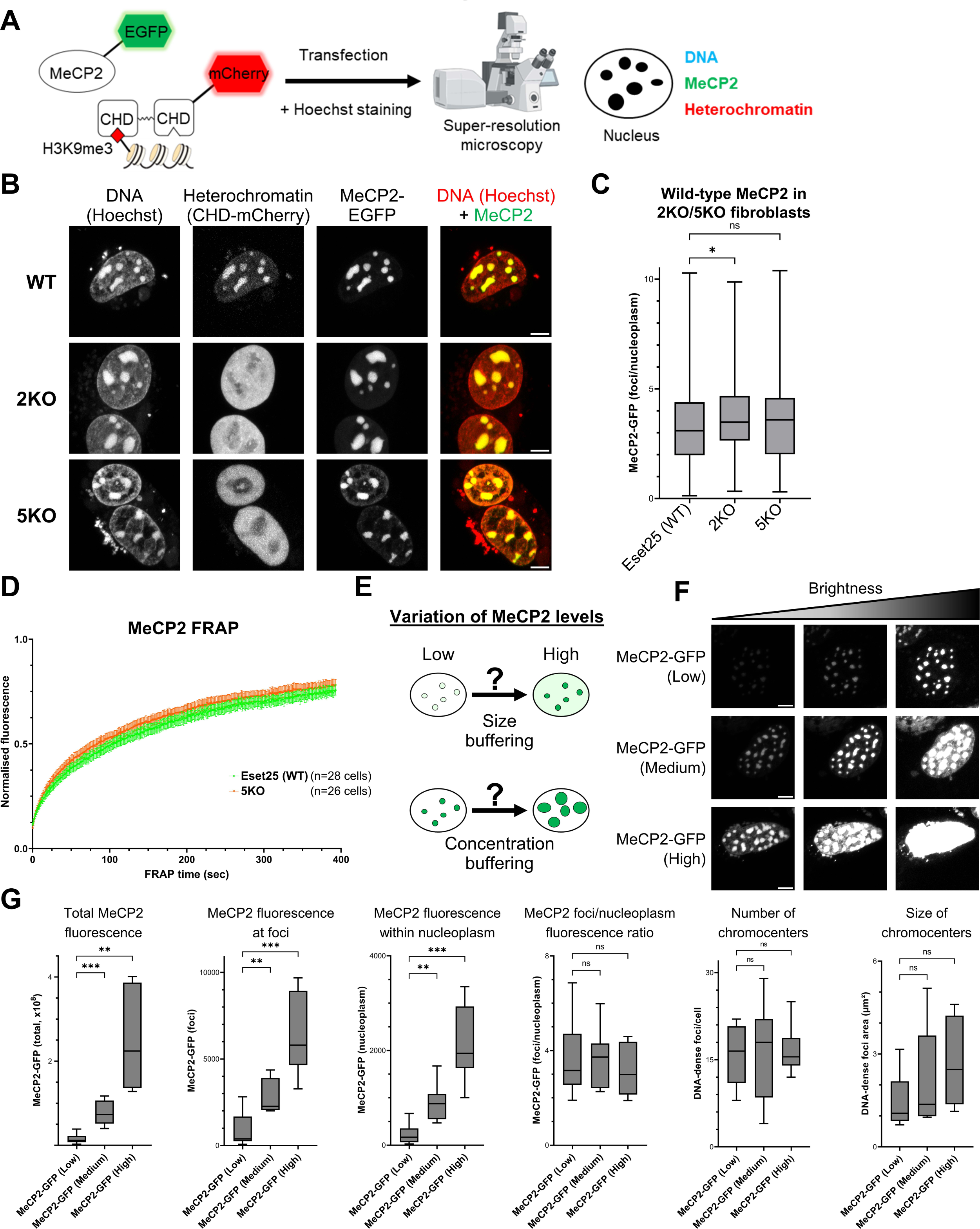
Heterochromatin organisation does not determine MeCP2 localisation. **A.** Diagram showing the strategy used to visualise MeCP2 (EGFP fusion protein), heterochromatin (HP1 chromodomain reporter) and DNA (Hoechst staining) in live cells by super-resolution microscopy. **B.** Live-cell imaging of wild-type (Eset25) and H3K9 lysine methyltransferases knockout fibroblasts (*2KO*/*5KO*) transfected with wild-type EGFP-MeCP2. Hoechst staining and CHD-mCherry reporter were used to visualise DNA and heterochromatin, respectively. Scale bars: 5µm. **C.** Box plot showing the quantification of MeCP2 fluorescence at DNA-dense foci (relative to nucleoplasm) in wild-type (Eset25), *2KO* and *5KO* fibroblasts, as described in panel B. **D.** Graph showing the FRAP quantification of wild-type EGFP-MeCP2 in wild-type (Eset25) and *5KO* fibroblasts. Error bars: SEM. **E.** Diagram showing two possible responses of MeCP2-containing foci to variations in expression levels. **F.** Live-cell imaging of *Mecp2* knockout fibroblasts transfected with varying levels of wild-type EGFP-MeCP2. Cells were divided into three expression categories (low, medium, high) and images are shown at three levels of brightness to enable comparison. Scale bars: 5µm. **G.** Box plots showing the quantification of different parameters in cells expressing varying levels of MeCP2, as described in panel F.

Transfection experiments allowed us to test further predictions of the hypothesis that MeCP2 modulates the structure of heterochromatic foci via its involvement in liquid-liquid phase separation (Brero *et al*, 2005; Bertulat *et al*, 2012; Li *et al*, 2020; Zhang *et al*, 2022).

This model predicts that increasing MeCP2 expression will result in larger chromocenters while maintaining a fixed concentration of MeCP2 (c_sat_) within each focus (Figure 2E); a property defined as “concentration buffering” (Banani *et al*, 2017). To test this, we analysed transfected *Mecp2*-null cells expressing varying levels of MeCP2 and found that the concentration of MeCP2 within foci did not remain constant but correlated with total MeCP2 expression levels (Figure 2F, 2G, S5D). As expected from a “size buffering” model (Figure 2E), the increase in MeCP2 within foci was also accompanied by increased MeCP2 concentration within the nucleoplasm, resulting in a relatively stable foci/nucleoplasm ratio of fluorescence signal (Figure 2G). Moreover, neither the size nor the number of heterochromatic foci was influenced by changes in MeCP2 expression (Figure 2G, S5D). Overall, these initial observations fail to support the phase separation model.

### MeCP2 nuclear distribution is diffuse in most mammalian species

A prediction of the phase separation hypothesis is that MeCP2, whose amino acid sequence is highly conserved in vertebrates, will form equivalent nuclear foci in cells from diverse mammalian species. To test this, we obtained cell lines (mainly primary fibroblasts) from 15 different species covering most mammalian lineages (Figure 3A) and transfected each with MeCP2-EGFP and CHD-mCherry. Surprisingly, super-resolution microscopy revealed a uniformly diffuse nuclear distribution of MeCP2 in the great majority of cell lines, including human. Furthermore, localisation of the heterochromatin reporter was dispersed throughout the nucleus (Figure 3B, 3C and S6). The only exceptions were mouse and red deer cells, where MeCP2 and heterochromatin co-localised within prominent foci. For unknown reasons, foci in red deer cells were only detected in about half of nuclei, the remainder showing a diffuse distribution. In addition, chromocenters characterised by intense foci of Hoechst staining in live cells were absent in most mammalian species, except mouse and red deer cells (Figure 3B, S6). The results indicate that MeCP2 does not have an intrinsic tendency to adopt a focal organisation.

**Figure 3.**
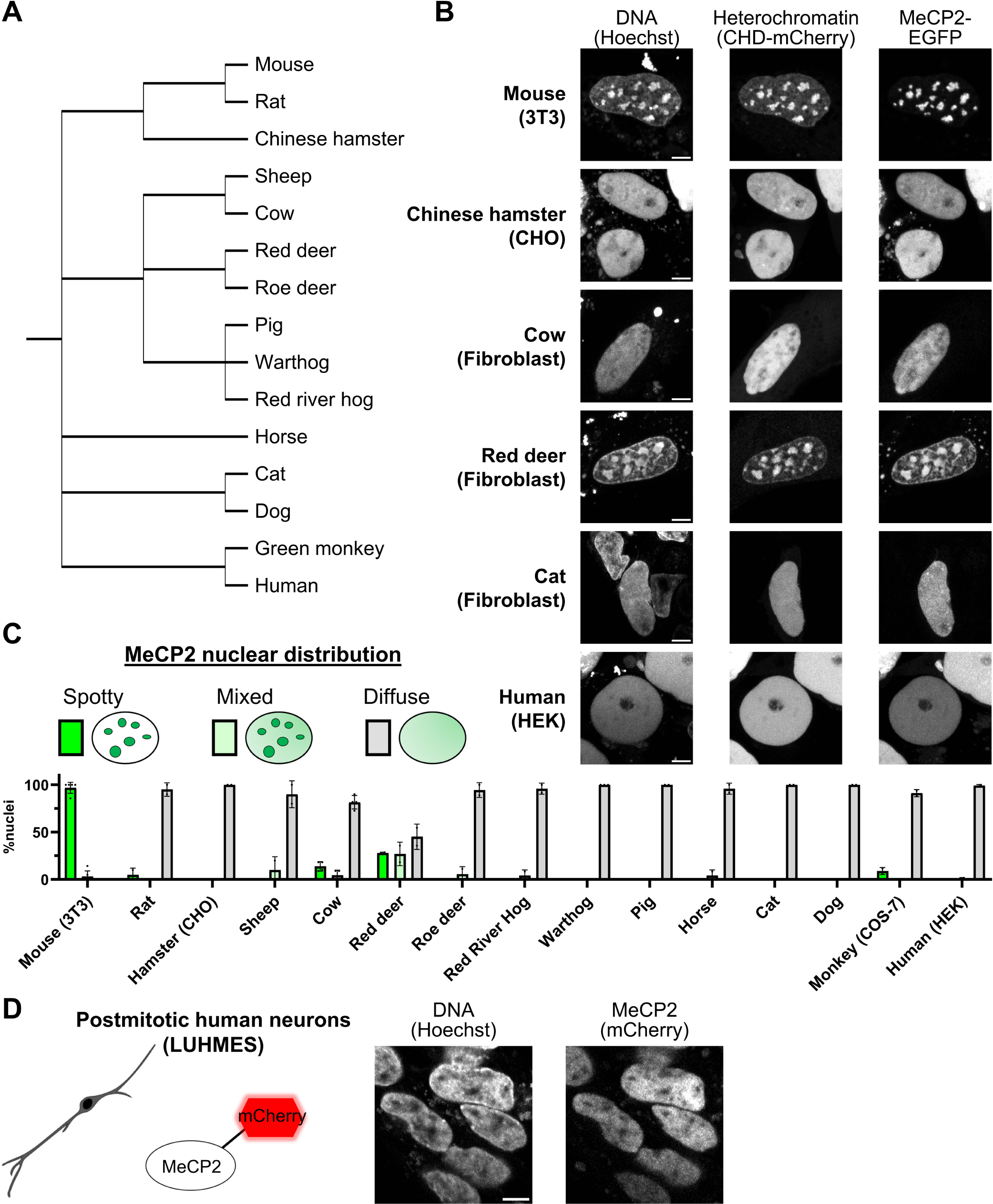
MeCP2 displays a diffuse nuclear distribution in most mammalian species. **A.** Phylogenetic tree showing the mammalian species used in this study (from NCBI Taxonomy). **B.** Live-cell imaging of the indicated mammalian cell lines transfected with wild-type EGFP-MeCP2. Hoechst staining and CHD-mCherry reporter were used to visualise DNA and heterochromatin, respectively. Scale bars: 5µm. **C.** Graph showing quantification of MeCP2 nuclear distribution (spotty, mixed, diffuse) in all studied mammalian species. **D.** Live-cell imaging of human postmitotic neurons (LUHMES cells) expressing endogenously tagged MeCP2-mCherry. Hoechst staining was used to visualise DNA. Scale bar: 5µm.

MeCP2 of mouse was expressed in all transfection experiments as the MeCP2 protein sequence is highly conserved across mammals (Figure S7). Notably, all residues within the MBD are identical in all the species analysed in this study. To test whether the observed nuclear distributions could nevertheless be due to the non-homologous origin of MeCP2, we transfected human MeCP2 into mouse and human fibroblasts. Again prominent nuclear foci were observed in mouse cells but MeCP2 was dispersed in human nuclei (Figure S8A).

Since MeCP2 is most abundant and functionally important in the nervous system, we also asked whether MeCP2 adopts a distinct distribution in neurons. We used human dopaminergic neurons derived from Lund Human Mesencephalic (LUHMES) cells (Lotharius *et al*, 2002; Scholz *et al*, 2011) that express endogenously tagged MeCP2-mCherry protein (Shah *et al*, 2016) (Figure S8B). As in fibroblasts, live-cell imaging confirmed that human neurons show a diffuse nuclear pattern of MeCP2 (Figure 3D). These findings indicate that the dramatic differences in MeCP2 nuclear localisation among mammals are not due to interspecific differences in their MeCP2 proteins or to the tissue origin of the cell type under investigation but are determined by distinctive genomic structures in the different species.

DNA methylation patterns are a prime candidate for determining whether MeCP2 is dispersed or concentrated in the nucleus (Figure 3C). Accordingly, immunostaining with an antibody directed against 5-methylcytosine showed large clusters resembling MeCP2 foci in mouse and red deer cells, while other species displayed diffuse patterns (Figure S9A). This observation is in agreement with the well-documented accumulation of methylated major satellite DNA at pericentric heterochromatin in mouse cells (Jones, 1970; Pardue & Gall, 1970; Rae & Franke, 1972; Miller *et al*, 1974; Manuelidis *et al*, 1982; Manuelidis, 1984; Guenatri *et al*, 2004). Like mouse, the red deer genome contains highly repetitive elements, including “satellite I” DNA which localises to pericentromeric regions (Lee & Lin, 1996; Lee *et al*, 1997; Vozdova *et al*, 2020). To test whether these satellite repeat elements are methylated in the red deer, we performed a Southern blot of genomic DNA digested with methylation-sensitive (HpaII) or methylation-insensitive (MspI) isoschizomers (Figure S9B). Probing with the satellite I DNA sequence confirmed that the repeat arrays are highly methylated (Figure S9C). Agarose gel electrophoresis also confirmed that satellite I DNA repeats are highly abundant, as they were visible as discrete bands in bulk genomic DNA by ethidium bromide staining. The results suggest that the demarcation of prominent chromocenters is dependent on reiteration of underlying satellite repeat DNA arrays and the presence of DNA methylation within the repeated motifs is the key genomic feature that leads to MeCP2 foci. To further explore this hypothesis, we compared fibroblast cell lines from two closely related mouse species that have dramatically different amounts of satellite repeat DNA: *Mus musculus*, the most widely used mouse model, and *Mus spretus* which diverged from *M. musculus* 1-2 million years ago (Staelens *et al*, 2002; Mahieu *et al*, 2006). In contrast to *Mus musculus* cells, the *Mus spretus* genome lacks almost all major satellite DNA repeats at pericentric regions (Brown & Dover, 1980; Matsuda & Chapman, 1991) (Figure 4A), as confirmed by Southern blot analysis (Figure 4B). Most major satellite repeats contain cleavage sites for both ApoI and HpyCH4IV (Figure S9D), but the latter enzyme cuts rarely due to methylation within its ACGT recognition sequence. Accordingly, immunostaining showed that 5-methycytosine is highly enriched at *Mus musculus* nuclear foci but appeared relatively diffuse in *Mus spretus* cells (Figure 4C). Live-cell imaging confirmed the absence of dense Hoechst-stained DNA foci, as well as diffuse patterns of MeCP2 and heterochromatic markers (Figure 4D, 4E). Overall, we conclude that the two exceptional species in the panel of mammals, mouse and red deer, differ from the majority by having satellite repeat DNA sequences near centromeres that are both highly abundant and methylated (Figure 4F).

**Figure 4.**
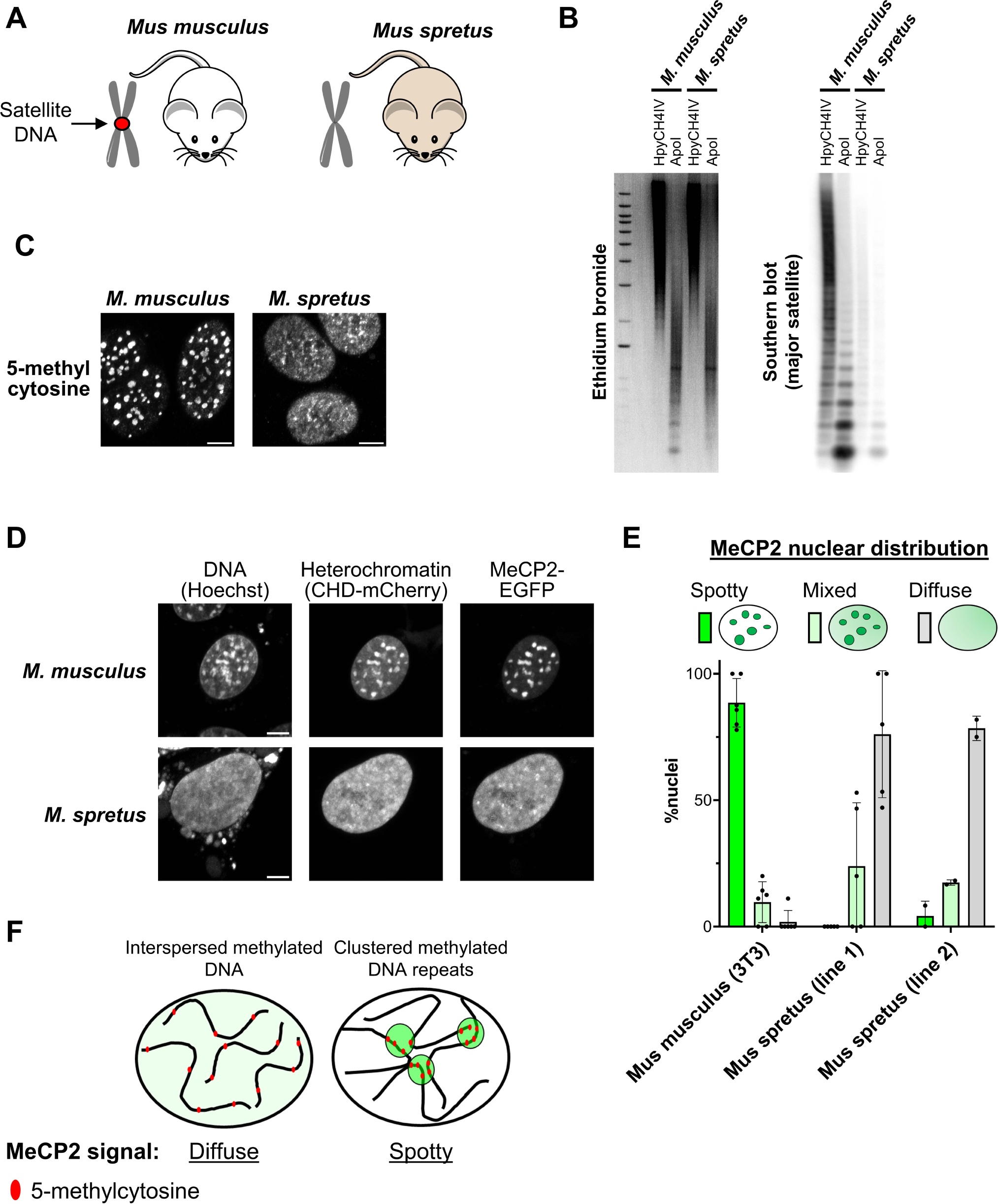
MeCP2 foci depend on the presence of abundant satellite DNA repeats. **A.** Diagram showing two closely related mouse strains used in this study. The *Mus musculus* genome contains abundant major satellite DNA repeats which cluster at pericentromeric regions, while *Mus spretus* lacks these repetitive elements. **B.** Ethidium bromide staining (left) and Southern blot (right) using a probe for major satellite DNA with *Mus musculus* and *Mus spretus* genomic DNA digested with a methylation-sensitive (HpyCH4IV) or -insensitive (ApoI) restriction enzymes. **C.** Immunofluorescence of antibody stained 5-methylcytosine in *Mus musculus* (3T3) and *Mus spretus* cell lines. Scale bars: 5µm. **D.** Live-cell imaging of *Mus musculus* (3T3) and *Mus spretus* cell lines transfected with wild-type EGFP-MeCP2. Hoechst staining and CHD-mCherry reporter were used to visualise DNA and heterochromatin, respectively. Scale bars: 5µm. **E.** Graph showing the quantification of MeCP2 nuclear distribution (spotty, mixed, diffuse) in *Mus musculus* (3T3) and *Mus spretus* cell lines. **F.** Model for differential MeCP2 nuclear distribution in mammalian species. In most species, including human, methylated DNA elements are interspersed along the genome leading to a diffuse MeCP2 nuclear pattern. In some cases, like in *Mus musculus*, methylated DNA repeats cluster into “chromocenters” leading to spotty MeCP2 signal.

## Discussion

The goal of this study was to unravel the relationship between MeCP2 accumulation, DNA methylation and heterochromatin organisation. In particular, we wished to assess using live-cell imaging the validity of recent claims that MeCP2 contributes to heterochromatin formation by undergoing phase separation. We found that the prominent MeCP2-containing foci seen in mouse cells are absent in 14 out of 16 mammalian species. Typically, the nuclear distribution of MeCP2 is diffuse in mammalian cells, including human neurons, indicating that MeCP2 does not intrinsically form large condensates. While purified MeCP2 protein forms liquid droplets *in vitro*, it fails to display key hallmarks of liquid-liquid phase separation *in vivo* (Banani *et al*, 2017; Alberti *et al*, 2019). Notably, in mouse cells we found no evidence for a critical concentration (c_sat_) at which MeCP2 would form condensates and showed that the size of MeCP2-containing foci is uncoupled from total MeCP2 expression levels. Our conclusions agree with a previous study that used live imaging approaches to show that MeCP2 does not behave as a member of a phase-separated compartment (Erdel *et al*, 2020).

The results also question the notion that mouse pericentric heterochromatin depends upon phase separation, as a heterochromatin reporter (Villaseñor *et al*, 2020) that recognises the histone mark H3K9me3 via its chromodomains revealed a diffuse nuclear pattern in most mammalian species. While heterochromatin is usually thought to involve chromatin compaction, most mammals in fact show little evidence of H3K9me3 clustering. These findings cast doubt on the proposal that heterochromatin-associated proteins, including HP1, drive heterochromatin condensation (Larson *et al*, 2017; Strom *et al*, 2017). As previously proposed, heterochromatic foci in mouse cells may resemble “collapsed chromatin globules” rather than phase-separated condensates (Erdel *et al*, 2020). The coalescence of pericentric repetitive DNA may result from structural features of satellite sequences, perhaps involving symmetrical DNA motifs that promote alternative conformations (Kasinathan & Henikoff, 2018). The AT-rich base composition of mouse major satellite is probably not required for condensation as satellite I of red deer is markedly more GC-rich that bulk genomic DNA (55% versus ∼40% GC).

Our data are compatible with a simple biochemical explanation for nuclear localisation of MeCP2 based on the affinity of MeCP2 for methylated DNA (Nan *et al*, 1996; Kudo *et al*, 2003; Marchi *et al*, 2007; Kumar *et al*, 2008; Schmiedeberg *et al*, 2009; Piccolo *et al*, 2019; Musacchio, 2022). The 85 amino acid MBD of MeCP2 alone is sufficient to target DNA-dense foci in mouse cells, but only in the presence of DNA methylation. Conversely, mutation of the MBD within the full-length protein abolishes MeCP2 subnuclear localisation in mouse cells. Although the AT-Hooks of MeCP2 may contribute to MeCP2 binding dynamics via weak and transient interactions with genomic DNA (Piccolo *et al*, 2019), these motifs are dispensable for MeCP2 localisation to heterochromatic foci. Moreover, mutations that inactivate the AT hook in humans apparently do not cause Rett syndrome (Lyst *et al*, 2016). Similarly, with the exception of the NCoR-interaction domain, intrinsically disordered regions are largely dispensable for MeCP2 function, as transgenic mice expressing a truncation of most N- and C-terminal IDRs show no overt phenotype (Tillotson *et al*, 2017). As previously noted (Fan *et al*, 2020), MeCP2 retains its localisation to DNA-dense foci in mouse cells in the complete absence of DNA methylation but its exchange as measured by FRAP is greatly increased, compatible with reduced DNA binding affinity. It is notable that mutations that similarly reduce the residence time of MeCP2 on mouse heterochromatin cause Rett syndrome (Schmiedeberg *et al*, 2009), suggesting that fast exchange is incompatible with MeCP2 function.

The nuclear distribution of MeCP2 appears to be directly influenced by variations in genomic DNA methylation patterns between species. In *Mus musculus* and red deer, MeCP2 coincides with dense clusters of 5-methylcytosine at arrays of satellite repeat sequences, whereas in the other 14 mammalian species tested, both DNA methylation and MeCP2 are relatively uniformly distributed, consistent with the global distribution of methylated CGs (Rabinowicz *et al*, 2003; Suzuki & Bird, 2008; de Mendoza *et al*, 2021; Klughammer *et al*, 2023). The importance of abundant repetitive DNA elements for the creation of MeCP2 foci is illustrated by the differences between two mouse species that are sufficiently closely related to form hybrids: *Mus musculus* and *Mus spretus*. *Mus spretus* cells lack most major satellite DNA repeats (Brown & Dover, 1980; Matsuda & Chapman, 1991), have no apparent chromocenters and, in striking contrast to the “classic” mouse model (*Mus musculus*), display diffuse nuclear MeCP2 and DNA methylation patterns. It should be noted that methylation of sequences other than CG, which is abundant in mature mammalian neurons and serves as an additional target for MeCP2 binding, is probably very rare in the cell lines under study. Accumulation of non-CG methylation to levels comparable with mCG occurs in mature neurons but is absent in non-neuronal cells and in neurons differentiated *ex vivo* (Lister *et al*, 2013; Guo *et al*, 2014; Cholewa-Waclaw *et al*, 2019).

Although MeCP2 localises prominently to heterochromatic foci in mouse, the protein is also bound to euchromatin genome wide. While the relationship between MeCP2 binding to euchromatic genes and transcriptional regulation has been extensively characterised (Gabel *et al*, 2015; Chen *et al*, 2015; Kinde *et al*, 2016; Lagger *et al*, 2017; Cholewa-Waclaw *et al*, 2019; Boxer *et al*, 2020; Tillotson *et al*, 2021), the functional significance of MeCP2 at pericentric heterochromatin remains unclear. Previous studies using cellular models proposed a role for MeCP2 in promoting the clustering of chromocenters (Brero *et al*, 2005; Marchi *et al*, 2007; Agarwal *et al*, 2011; Bertulat *et al*, 2012) or by controlling the partitioning of HP1(α/γ) within foci (Agarwal *et al*, 2007; Li *et al*, 2020). Additionally, it was reported that MeCP2 directly recruits H3K9 methyltransferase activity (Fuks *et al*, 2003). However, *MeCP2*-null neurons in mice show patterns of H3K9me3 and HP1 in heterochromatin which are largely indistinguishable from wild-type, although slightly increased DAPI staining within foci and altered patterns of ATRX or H4K20me3 have been reported (Nan *et al*, 2007; Linhoff *et al*, 2015; Ito-Ishida *et al*, 2020). Moreover, complete loss of H3K9 methylation and consequent dispersion of HP1α does not affect MeCP2 localisation to foci in mouse cells and has no impact on its DNA binding dynamics as assessed by FRAP. Thus, taken together, the results presented here argue that the biogenesis of heterochromatin compartments does not involve or require MeCP2.

## Acknowledgements

We thank Verdiana Steccanella and Jenna Hare for technical assistance. We also thank Gura Bergkvist, Xavier Donadeu, Bruce Whitelaw, Jacky Guy and Josephine Pemberton (University of Edinburgh) for sharing mammalian cell lines and genomic DNA used in this study, and Tuncay Baubec (Utrecht University) for sharing the chromodomain reporter construct. Imaging was performed in Centre Optical Instrumentation Laboratory supported by a Core Grant (203149) to the Wellcome Centre for Cell Biology at the University of Edinburgh. This work was funded by European Research Council Advanced Grant EC 694295 Gen-Epix and Wellcome Investigator Award #107930. A.B. is a member and K.P. is supported by the Simons Initiative for the Developing Brain grant (SFARI-529085). The collection of primary cell lines from mammalian species was funded by a BBSRC Institute Strategic Programme grant BBS/E/D/10002071 to T.B. Research in the laboratory of T.J. is supported by the Max Planck Society and by grants (CRC992 “MEDEP”) from the German Research Foundation (DFG).

## Author contributions

Conceptualization: A.B., R.P.; Methodology: R.P., M.B., S.H., K.P., T.B., S.M.; Software: T.M.2., D.A.K.; Validation: R.P., M.B., S.H.; Formal analysis: R.P.; Investigation: R.P., M.B., S.H., K.P.; Resources: T.B., S.M., T.M.1., T.J., T.H., C.L.; Writing - Original Draft: A.B., R.P.; Writing - Review & Editing: R.P., M.B., K.P., T.B., T.J., T.M.2., A.B.; Supervision: A.B., R.P.; Funding acquisition: A.B., T.B., T.J. T.M.1.: Thomas Montavon; T.M.2.: Toni McHugh

## Materials and methods

### Cell culture

All cell lines were grown at 37°C with 5% CO_2_. Fibroblasts and immortalised cell lines were cultured in Glasgow minimum essential medium (GMEM; Gibco ref. 11710035) supplemented with 10% fetal bovine serum (batch tested), 1x L-glutamine (Gibco ref. 25030024), 1x MEM non-essential amino acids (Gibco ref.11140035), 1mM sodium pyruvate (Gibco ref. 11360039), 0.1mM 2-mercaptoethanol (Gibco ref. 31350010). Eset25, 2KO and 5KO cell lines were grown in Dulbecco’s Modified Eagle Medium (DMEM, Gibco ref. 41966029) supplemented with 10% fetal bovine serum and 1.25µg/ml puromycin. ESC lines were cultured in gelatin coated dishes containing GMEM supplemented with 10% fetal bovine serum, 1x L-glutamine, 1x MEM non-essential amino acids, 1mM sodium pyruvate, 0.1mM 2-mercaptoethanol and 100U/ml leukemia inhibitory factor (LIF, batch tested).

LUHMES cells were cultured and differentiated as described (Scholz *et al*, 2011; Shah *et al*, 2016) in poly-L-ornithine and fibronectin coated dishes. Primary cell lines were derived from tissue explants or collagenase dissociated tissues and expanded in GMEM supplemented with 10% fetal bovine serum, 2mM L-glutamine, 1x MEM non-essential amino acids, 1mM sodium pyruvate, 0.1mM 2-mercaptoethanol, 100U/ml Penicillin-Streptomycin, 50µg/ml Gentamicin and 2.5µg/ml Amphotericin B. Once established, cultures were maintained in the same medium without antibiotics. *Mus spretus* primary fibroblasts were immortalised using SV40 virus as described (Hendrich *et al*, 2001). For imaging, cells were plated and cultured directly on polymer coverslips (iBidi cat. 81156 or 80286) with the appropriate coating and culture conditions.

As a quality control, all mammalian cell lines were tested for Mycoplasma contamination (Lonza cat. LT07-218). Additionally, the identity of each cell line was verified by Sanger sequencing of mitochondrial Cytochrome b which was amplified from extracted total DNA using universal primers (FW: CGAAGCTTGATATGAAAAACCATCGTTG, RV: AAACTGCAGCCCCTCAGAATGATATTTGTCCTCA) (Kocher *et al*, 1989; Irwin *et al*, 1991).

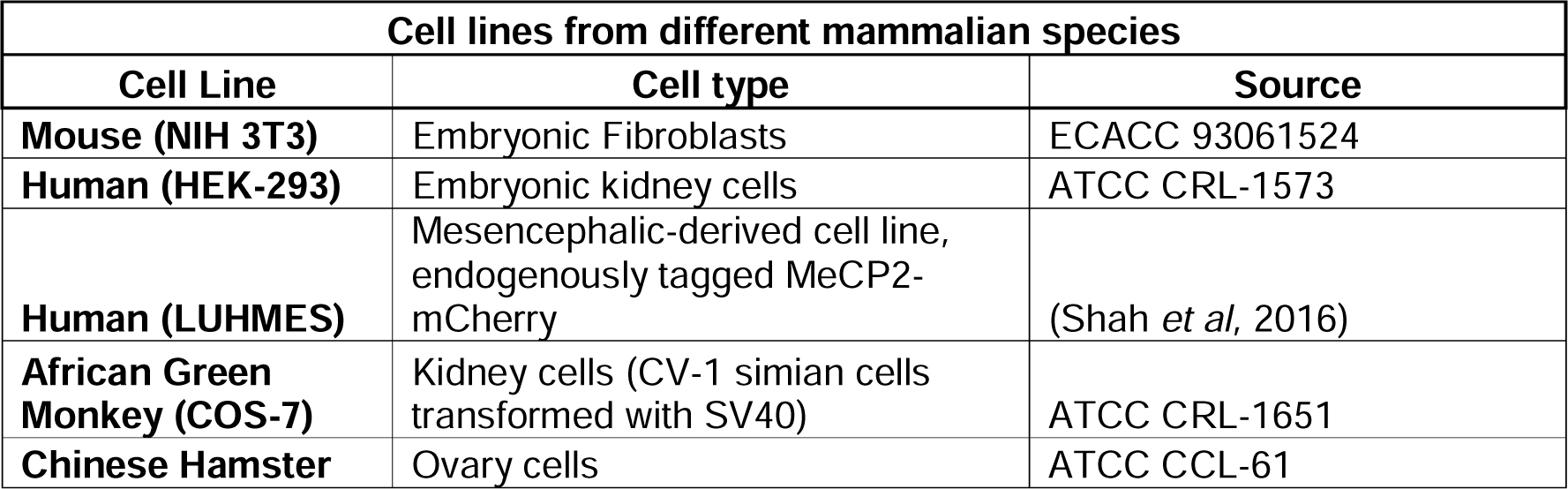
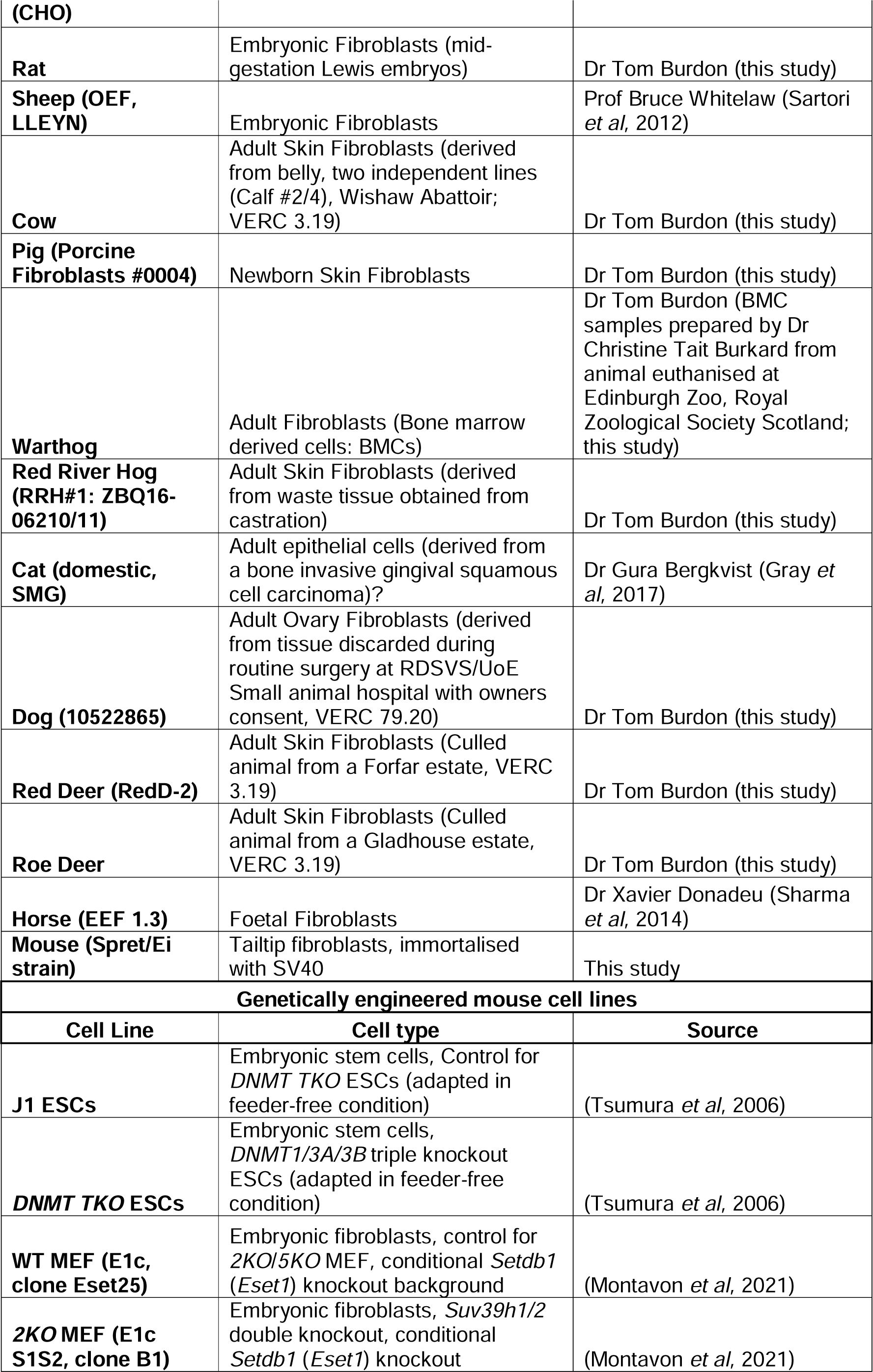
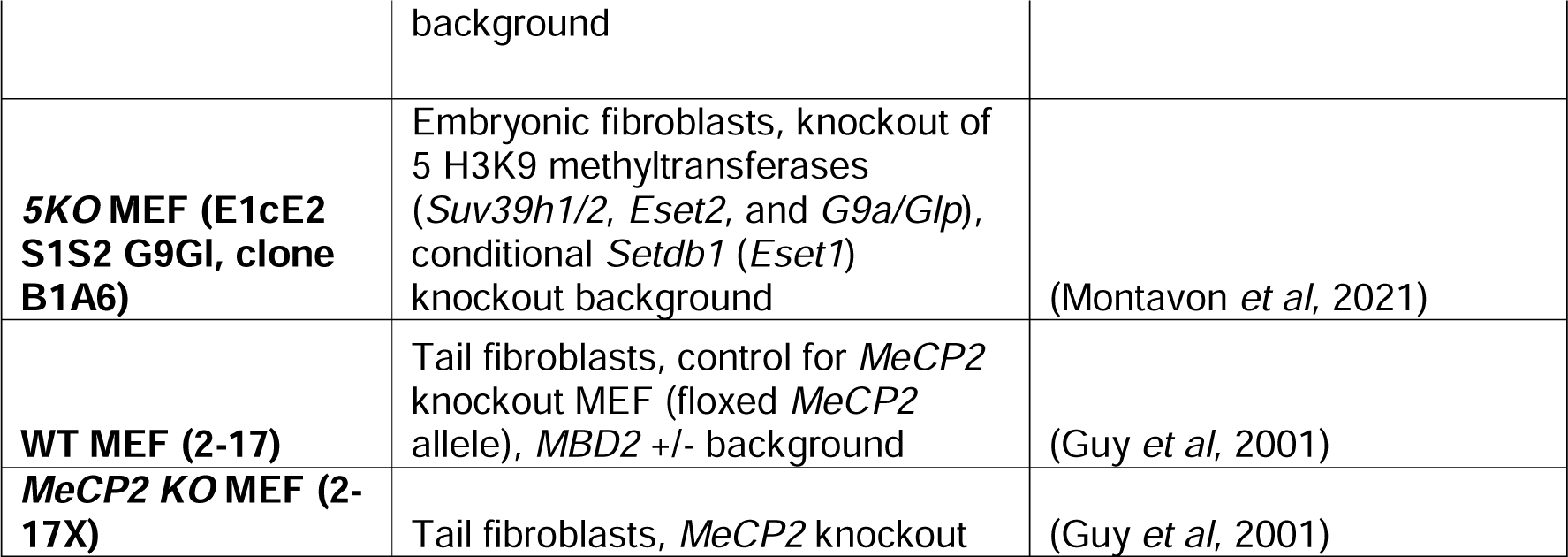

### Molecular cloning

For the heterochromatin reporter, the dimerised CBX1 chromodomain from a previously described engineered chromatin reader (eCR) (Villaseñor *et al*, 2020) was subcloned into a pPYCAG mammalian expression vector (Chambers *et al*, 2003) containing mCherry. MeCP2 mutant constructs were cloned by Gibson assembly (NEB cat. E5520S) using synthetic double-stranded DNA fragments ordered from Integrated DNA Technologies (IDT).

| Mammalian expression plasmids |  |  |
| --- | --- | --- |
| Name | Details | Source |
| pPYCAG-mCherry-2xCHD | Heterochromatin reporter (dimerised CBX1 chromodomain), N-terminal mCherry tag, Constitutive CAG enhancer/promoter | This study |
| pEGFP-Human MeCP2 | Wild-type Human MeCP2, N-terminal EGFP tag, Constitutive CMV enhancer/promoter | (Klose *et al*, 2005) |
| pEGFP-Mouse MeCP2 | Wild-type Mouse MeCP2, N-terminal EGFP tag, Constitutive CMV enhancer/promoter | (Guy *et al*, 2018) |
| pEGFP-MeCP2 ATHookmut | Mouse MeCP2 AT Hook mutant (R188G, R190G), N-terminal EGFP tag, Constitutive CMV enhancer/promoter | This study |
| pEGFP-MeCP2 ΔMBD | Mouse MeCP2 MBD mutant (Δ99-125), N-terminal EGFP tag, Constitutive CMV enhancer/promoter | This study |
| pEGFP-MeCP2 ΔMBD + ATHookmut | Mouse MeCP2 MBD and AT Hook double mutant (Δ99-125, R188G, R190G), N-terminal EGFP tag, Constitutive CMV enhancer/promoter | This study |
| pEGFP-MeCP2 Minimal MBD | Mouse Minimal functional MBD (78-162), N-terminal EGFP tag, Constitutive CMV enhancer/promoter | This study |

### Live-cell imaging

Cells were transfected with MeCP2-EGFP with or without CHD-mCherry plasmid using Lipofectamine 3000 (Thermo Fisher Scientific cat. L3000008) and following the manufacturer’s protocol. Cells were analysed the next day following a medium change and Hoechst 33342 (Thermo Fisher Scientific cat. R37605) staining using the Zeiss LSM 880 confocal microscope (with Airyscan module) at 37°C with 5% CO_2_. Images were processed using the software Fiji.

To quantify fluorescence at DNA-dense foci in mouse cells, images were processed using a custom script (https://doi.org/10.5281/zenodo.7740611). The detection of nuclei was automated by performing image segmentation on the Hoechst fluorescence channel (maximum intensity Z-projection) using the software Cellpose (Stringer *et al*, 2021) integrated into the script using the BIOP ijl-utilities-wrappers plugin (v0.3.19). DNA-dense foci (chromocenters) were detected by applying thresholds per nucleus and several parameters were measured including the area of foci and nuclei (µm²), fluorescence levels (average pixel intensity) of MeCP2/Hoechst at foci and within the nucleoplasm (total nuclear signal depleted from foci). For statistical comparisons between MeCP2 wild-type and mutant foci, we performed a Brown-Forsythe and Welch one-way ANOVA test using the Games-Howell’s method to correct for multiple comparisons (adjusted p value) using the software GraphPad Prism 9. For the quantification of MeCP2 nuclear distribution in mammalian species, all transfected cells were counted and classified into three categories (spotty, mixed, diffuse). Results were plotted using GraphPad Prism 9.

### FRAP

Cells were transfected with wild-type MeCP2-EGFP using Lipofectamine 3000 (Thermo Fisher Scientific cat. L3000008) and following manufacturer’s protocol. Cells were analysed the next day following a medium change, and FRAP was performed as previously described (Tillotson *et al*, 2021) using the Zeiss LSM 880 confocal microscope at 37°C with 5% CO_2_. For each cell, EGFP-MeCP2 signal was imaged every 1sec for 400sec with five images recorded before bleaching a selected MeCP2 spot (FRAP spot) with 100% laser power. Images were processed using the software Fiji.

FRAP analysis of two independent transfection experiments was performed as previously described (Tillotson *et al*, 2021) using a custom macro with Fiji software (https://doi.org/10.5281/zenodo.2654601). Fluorescence was measured at the bleached MeCP2 spot (FRAP spot), as well as a non-bleached MeCP2 spot (control spot) to account for photobleaching during the experiment. Additionally, the fluorescence outside of transfected cells was measured as background. The first time point (T_0_) was defined as the first post-bleach image. For each time point, the FRAP fluorescence signal was normalised to fluorescence values before photobleaching, as described in the equation below:

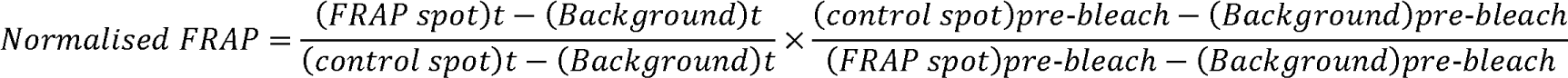

A “Two phase association” model (nonlinear regression) was used to fit experimental data using the software GraphPad Prism 9. The plateau (fluorescence at the last time point of the FRAP experiment), corresponding to the mobile fraction, was interpolated from the fitted curve. Conversely, the immobile fraction corresponds to the pool of fluorescent proteins not recovered during the experiment (=100%-mobile fraction). The T_50_ (time to recover 50% of pre-bleach level) was interpolated from the fitted curve. Data was plotted using GraphPad Prism 9.

### Immunofluorescence

Immunofluorescence was performed as previously described (Pantier *et al*, 2021). Cells were washed with PBS and fixed with 4% PFA for 10min at room temperature. After fixation, cells were washed with PBS and permeabilised for 10min at room temperature in PBS supplemented with 0.3% (v/v) Triton X-100. Samples were blocked for 1h30min at room temperature in blocking buffer: PBS supplemented with 0.1% (v/v) Triton X-100, 1% (w/v) BSA and 3% (v/v) serum of the same species as secondary antibodies were raised in (ordered from Sigma-Aldrich). Following blocking, samples were incubated overnight at 4°C with primary antibodies (see Table) diluted at the appropriate concentration in blocking buffer. After 4 washes in PBS supplemented with 0.1% (v/v) Triton X-100, samples were incubated for 2h at room temperature (in the dark) with fluorescently labelled secondary antibodies (Invitrogen Alexa Fluor Plus antibodies) diluted (1:500) in blocking buffer. Cells were washed 4 times with PBS supplemented with 0.1% (v/v) Triton X-100. DNA was stained with DAPI for 5min at room temperature, and cells were submitted to a final wash with PBS. Samples were imaged using the Zeiss LSM 880 confocal microscope (with Airyscan module). Images were processed using the software Fiji.

For 5mC immunostaining, we performed the same protocol as described above with extra steps. Following permeabilization, DNA was denatured with 4M HCl for 10min at 37°C. The pH of the medium was subsequently neutralised with three quick washes in PBS supplemented with 0.1% (v/v) Triton X-100. We then proceeded with the blocking step and followed the rest of the immunostaining protocol.

| Antibodies |  |  |
| --- | --- | --- |
| Target | Working dilution | Reference |
| **H3K9me3** (Rabbit polyclonal antibody) | 1:200 | Abcam cat. ab8898 |
| **HP1α** (Rabbit polyclonal antibody) | 1:200 | Cell Signaling cat. 2616 |
| **MeCP2** (Rabbit monoclonal antibody) | 1:200 | Cell Signaling cat. 3456 |
| **TUJ1 (TUBB3)** (Mouse monoclonal antibody) | 1:500 | BioLegend cat. 801201 |
| **5-methylcytosine** (Mouse monoclonal antibody) | 1:1,000 | Active Motif cat. 39649 |

### Southern blot

Genomic DNA was extracted using a commercial kit (Qiagen cat. 158388) and Southern blot was performed as previously described (Paton *et al*, 2022). Equal amounts of genomic DNA were digested and loaded into each lane. Mouse (*M. musculus* and *M. spretus*) genomic DNA was digested with methylation-sensitive (HpyCH4IV, NEB cat. R0619L) or methylation-insensitive (ApoI, NEB cat. R3566L) restriction enzymes cutting within major satellite DNA repeats. Red deer genomic DNA was digested with methylation-sensitive (HpaII, NEB cat. R0171M) or methylation-insensitive (MspI, NEB cat. R0106M) restriction enzymes recognising the same ‘CCGG’ sequence within satellite I DNA. For detection of mouse major satellite DNA, we used a 804bp probe including three repeats of the consensus sequence (Hörz & Altenburger, 1981), which was PCR-amplified from genomic DNA and cloned into a plasmid. For detection of red deer satellite DNA, we used a 725bp probe corresponding to the satellite I consensus sequence (GenBank accession numbers: U48429, MT185963, MT185964, MT185965, MT185966) (Lee & Lin, 1996; Vozdova *et al*, 2020), which was ordered as a synthetic double-stranded DNA fragment from Integrated DNA Technologies (IDT).

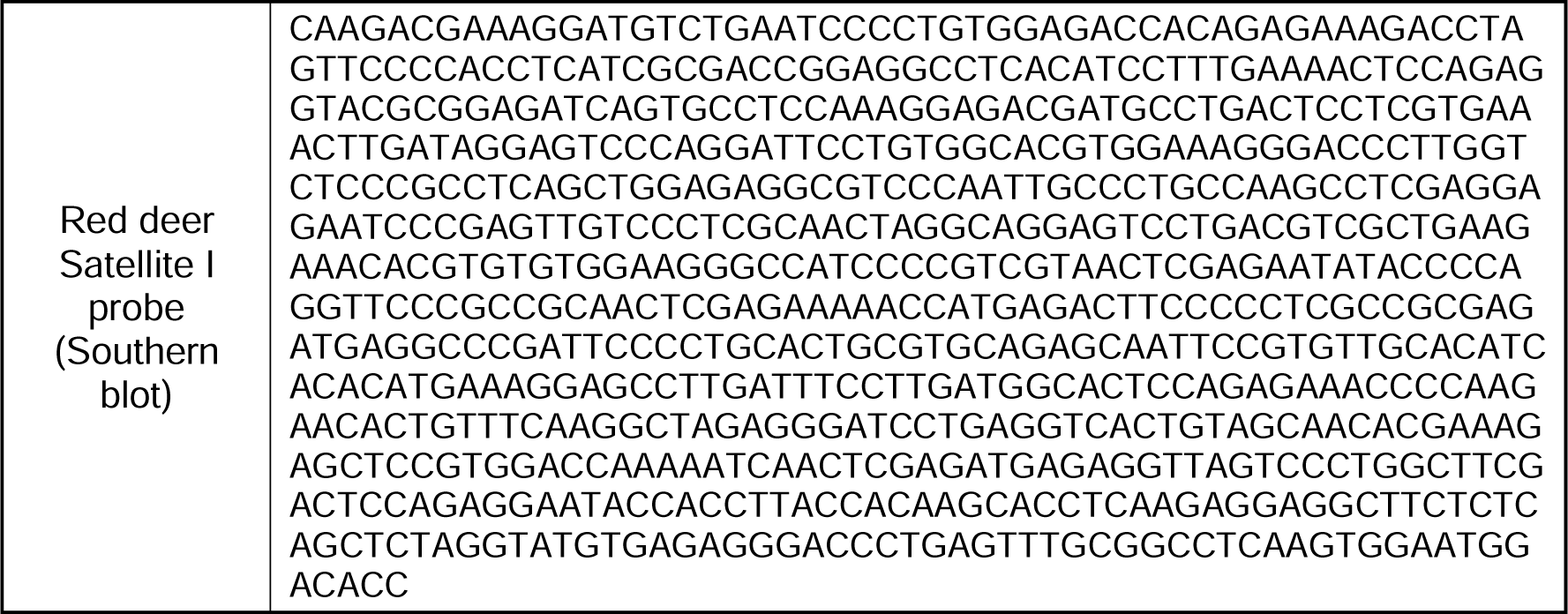
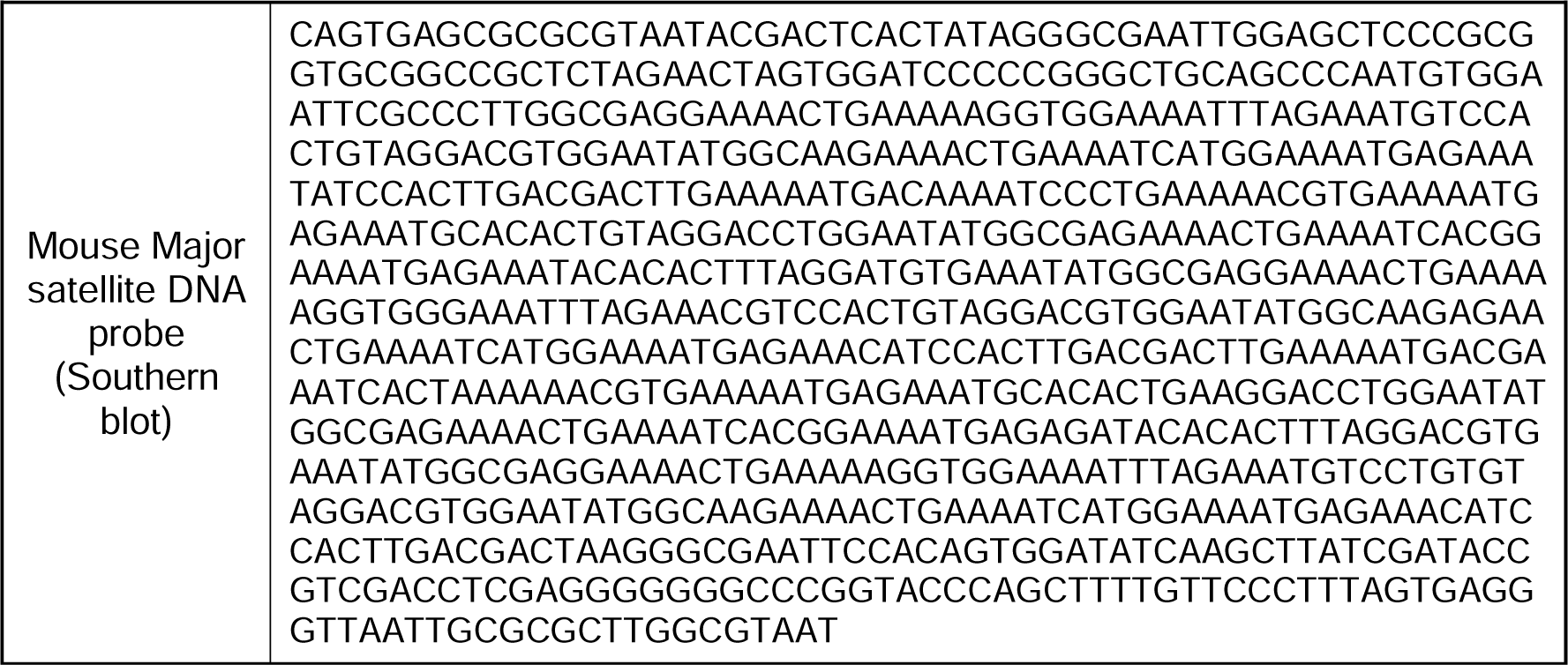

### 5mC LC-MS

Genomic DNA from two independent biological replicates was extracted using a commercial kit (Qiagen cat. 69504) with an optional RNase A treatment step. The quantification of global 5-methylcytosine levels was performed by Zymo Research (US) using an SRM-based mass spectrometry assay. Samples were measured in technical triplicates and data was represented as total amount of 5-methylcytosine relative to the total pool of guanine within genomic DNA (%mC/G). Data plotted on GraphPad Prism 9.

### Protein alignment

MeCP2 protein sequences of the mammalian species used in this study were obtained from the UniProt database (UniProt identifiers: Q9Z2D6 (Mouse), P51608 (Human), Q00566 (Rat), A0A3Q2HTZ1 (Horse), A0A8C2MED1 (Chinese hamster), M3WF10 (Cat), A0A8I3SAQ1 (Dog), A0A3Q1M211 (Cow), A0A8D1PU92 (Pig)). For other species for which a complete MeCP2 protein sequence was unavailable, we obtained transcript sequences (Ensembl ENSOART00020034580 (Sheep), NCBI XM_007993126 (African green monkey), NCBI XM_043895361 (Red deer)) and translated them *in silico*. All protein sequences were aligned using the Clustal Omega programme available as an online tool on the UniProt website (https://www.uniprot.org/align). Alignment data was visualised using the software ESPRIPT (https://espript.ibcp.fr/ESPript/ESPript/index.php) (Robert & Gouet, 2014).

### Data Availability

Raw and processed data used to generate the figures will be available on Mendeley/Zenodo (link). Reagents used in this study are available upon request.

## Supplementary figure legends

**Supplementary Figure 1 (related to Figure 1)**

A. Diagram showing the MeCP2 wild-type and mutant constructs (fused with EGFP) used for live-cell imaging. MBD: Methyl-DNA Binding Domain, NID: NCoR-Interaction Domain, IDR: Intrinsically Disordered Region. B. Live-cell imaging of 3T3 cells transfected with the EGFP-MeCP2 constructs indicated in panel A. Hoechst staining was used to visualise DNA. Scale bars: 5µm. C. Box plot showing the quantification of MeCP2 wild-type and mutants fluorescence at DNA-dense foci (relative to nucleoplasm) in 3T3 cells, as described in panel B.

**Supplementary Figure 2 (related to Figure 1)**

A. Live-cell imaging of J1 ESCs transfected with EGFP-MeCP2 wild-type and mutant constructs (see Figure S1). Hoechst staining was used to visualise DNA. Scale bars: 5µm. B. Box plot showing the quantification of wild-type and mutant MeCP2 fluorescence at DNA-dense foci (relative to nucleoplasm) in J1 ESCs, as described in panel A.

**Supplementary Figure 3 (related to Figure 1)**

A. Mass spectrometric quantification of 5-methylcytosine (relative to total guanosine pool) from genomic DNA in the indicated ESC lines. Individual data points are technical replicates. B. Live-cell imaging of *DNMT TKO* ESCs transfected with EGFP-MeCP2wild-type and mutant constructs (see Figure S1). Hoechst staining was used to visualise DNA. Scale bars: 5µm. C. Box plot showing the quantification of wild-type and mutant MeCP2fluorescence at DNA-dense foci (relative to nucleoplasm) in *DNMT TKO* ESCs, as described in panel B. D. Live-cell imaging showing the fluorescence recovery after photobleaching (FRAP) of wild-type EGFP-MeCP2 in wild-type (J1) and *DNMT TKO* ESCs. Scale bars: 5µm.

**Supplementary Figure 4 (related to Figure 2)**

**A, B**. Immunofluorescence of H3K9me3 (A) and HP1α (B) in wild-type (Eset25) and H3K9 lysine methyltransferases knockout fibroblasts (*2KO*/*5KO*). DAPI staining was used to visualise DNA. Scale bars: 5µm. **C**. Live-cell imaging of *5KO* fibroblasts transfected with EGFP-MeCP2 wild-type and mutant constructs (see Figure S1). Hoechst staining and CHD-mCherry reporter were used to visualise DNA and heterochromatin, respectively. Scale bars: 5µm. **D**. Box plot showing the quantification of MeCP2 wild-type and mutants fluorescence at DNA-dense foci (relative to nucleoplasm) in *5KO* fibroblasts, as described in panel C. **E**. Live-cell imaging showing the fluorescence recovery after photobleaching (FRAP) of wild-type EGFP-MeCP2 in wild-type (Eset25) and *5KO* fibroblasts. Scale bars: 5µm. **F**. Graph showing the FRAP quantification of wild-type EGFP-MeCP2 in all mutant lines used in this study (see Figures 1 and 2). Error bars: SEM.

**Supplementary Figure 5 (related to Figure 2)**

**A, B, C.** Immunofluorescence of MeCP2 (A), H3K9me3 (B) and HP1α (C) in wild-type (2-17) and *MeCP2* knockout fibroblasts. DAPI staining was used to visualise DNA. Scale bars: 5µm. D. Scatterplots showing the relationship between total MeCP2 expression levels within cells and different parameters associated with MeCP2 foci. The calculated R² values indicate a significant correlation only between total MeCP2 expression and MeCP2 concentration within foci.

**Supplementary Figure 6 (related to Figure 3)**

Live-cell imaging of all studied mammalian cell lines transfected with wild-type EGFP-MeCP2.Hoechst staining and CHD-mCherry reporter were used to visualise DNA and heterochromatin, respectively. Scale bars: 5µm. Of note, red deer cells presented two distinct phenotypes characterised by diffuse or spotty MeCP2/Heterochromatin, respectively.

**Supplementary Figure 7 (related to Figure 3)**

Protein alignment of MeCP2 across mammalian species. Identical residues are white on a red background; conservative substitutions found in several mammalian species are in red. Brackets indicate the MBD domain (residues 78-162) which is strictly conserved.

**Supplementary Figure 8 (related to Figure 3)**

A. Live-cell imaging of mouse (3T3) and human (HEK) cell lines transfected with mouse (left) or human (right) wild-type EGFP-MeCP2.Hoechst staining was used to visualise DNA. Scale bars: 5µm. B. Immunofluorescence of TUJ1in human postmitotic neurons (LUHMES) expressing endogenously tagged MeCP2-mCherry. DAPI staining was used to visualise DNA. Scale bar: 50µm.

**Supplementary Figure 9 (related to Figure 4)**

**A.** Immunofluorescence of 5-methylcytosine in the indicated mammalian cell lines. Scale bars: 5µm. **B.** Consensus sequence of red deer satellite I. The restriction site for HpaII/MspI enzymes used for Southern blot is highlighted in orange. **C.** Ethidium bromide staining (left) and Southern blot (right) using a probe for satellite I DNA repeats with red deer genomic DNA digested with a methylation-sensitive (HpaII) or -insensitive (MspI) restriction enzyme. Discrete bands at the bottom of the gel with MspI, but not HpaII, digested DNA indicates abundant and highly methylated DNA repeats in red deer cells. **D.** Consensus sequence of mouse major satellite DNA. The restriction sites for HpyCH4IV and ApoI enzymes used for Southern blot are highlighted in red and blue, respectively.

