## Supplementary Figures for "MeCP2 binds to methylated DNA independently of phase separation and heterochromatin organisation"

### Figure S1 (related to Figure 1) – 3T3 cells

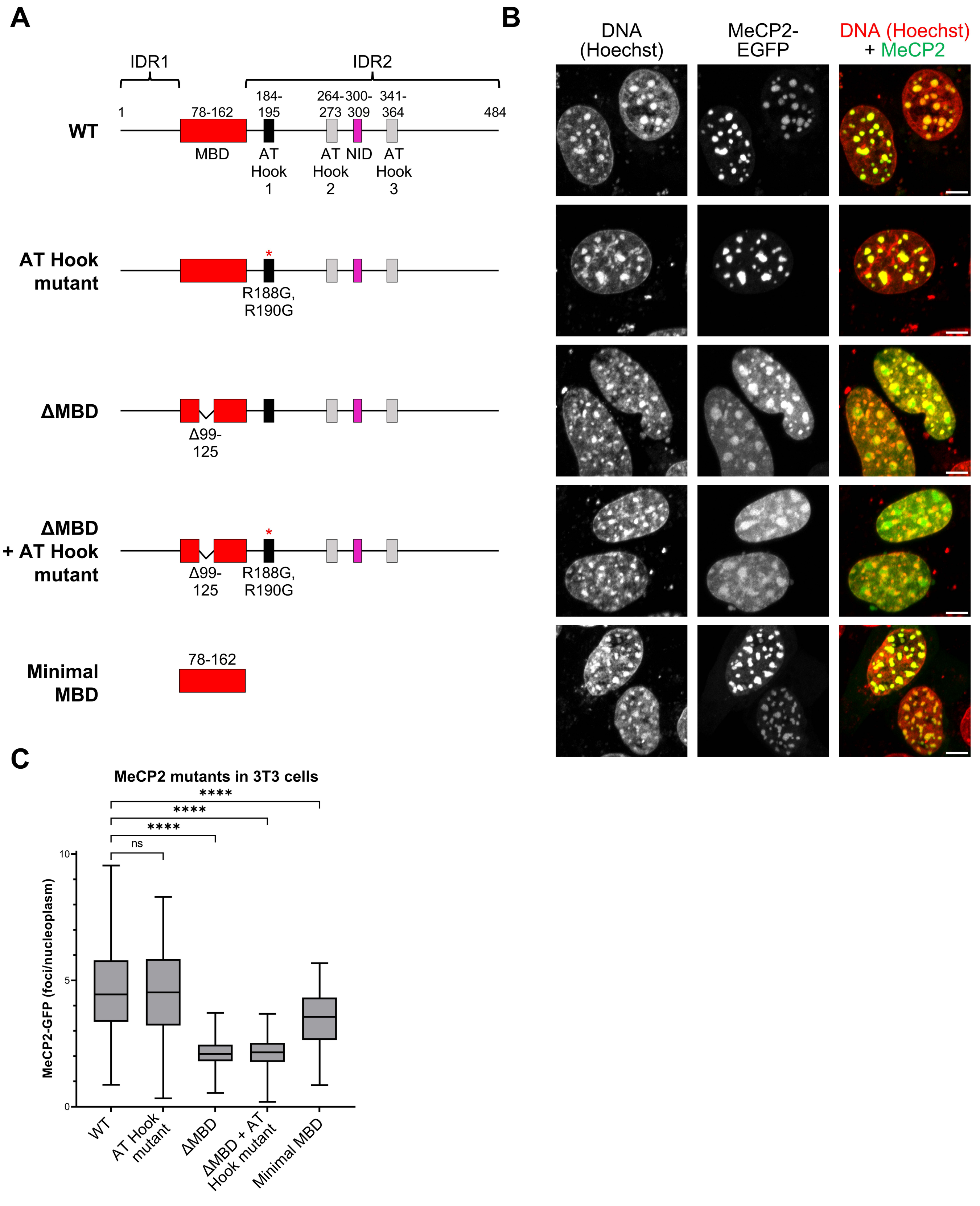

### Figure S2 (related to Figure 1) – J1 ESCs

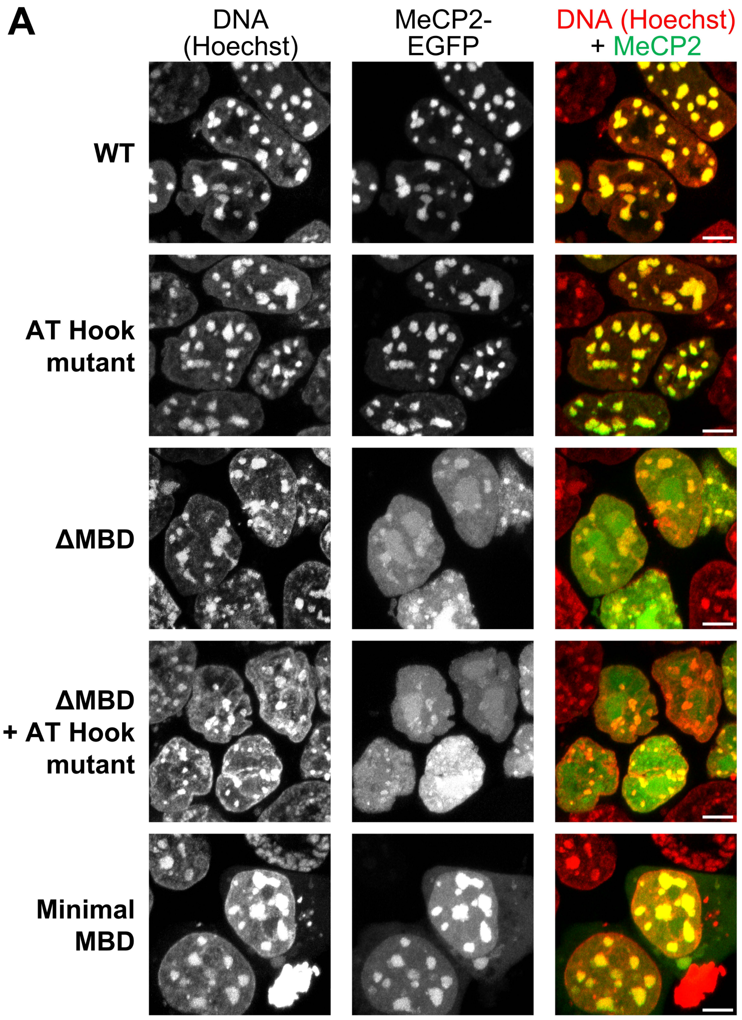


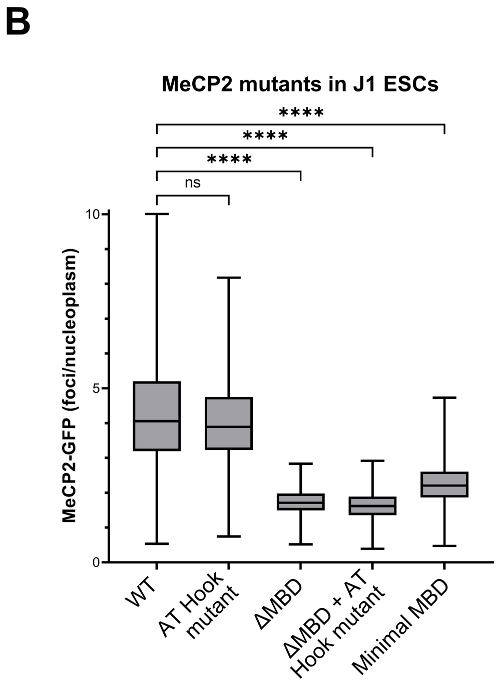

### Figure S3 (related to Figure 1) – DNMT TKO ESCs

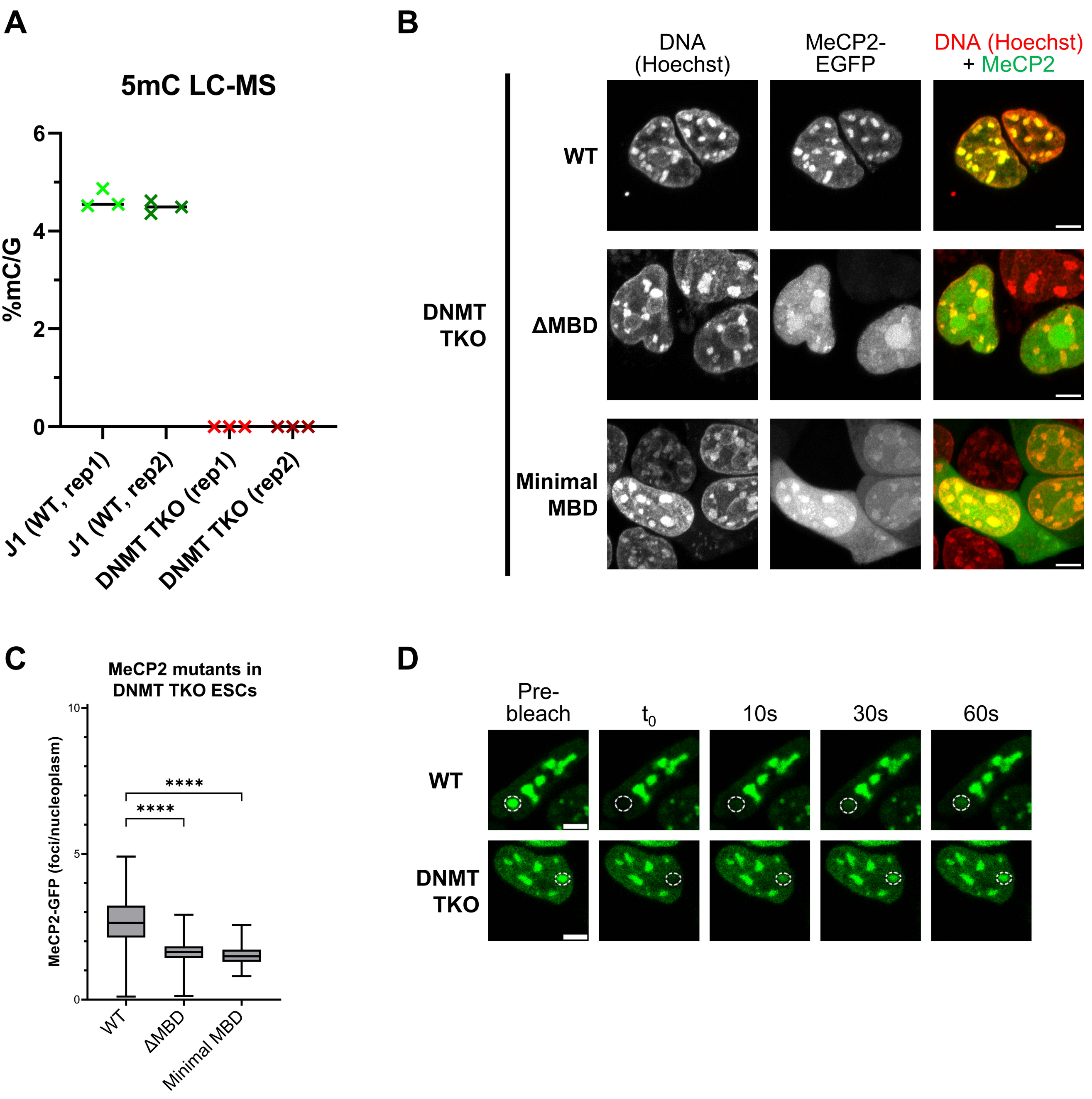

### Figure S4 (related to Figure 2) – 2KO/5KO cells

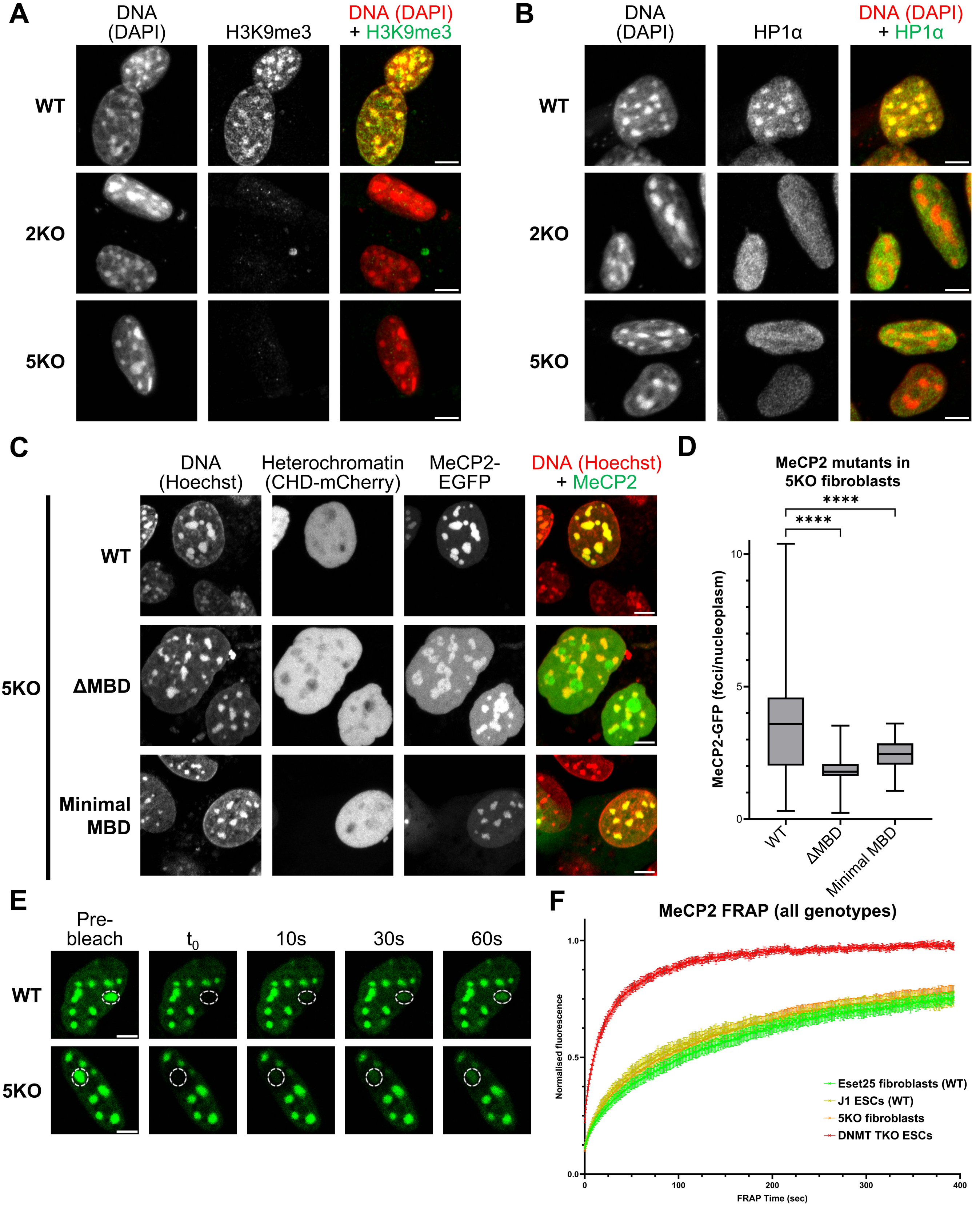

### Figure S5 (related to Figure 2) – MeCP2 KO cells

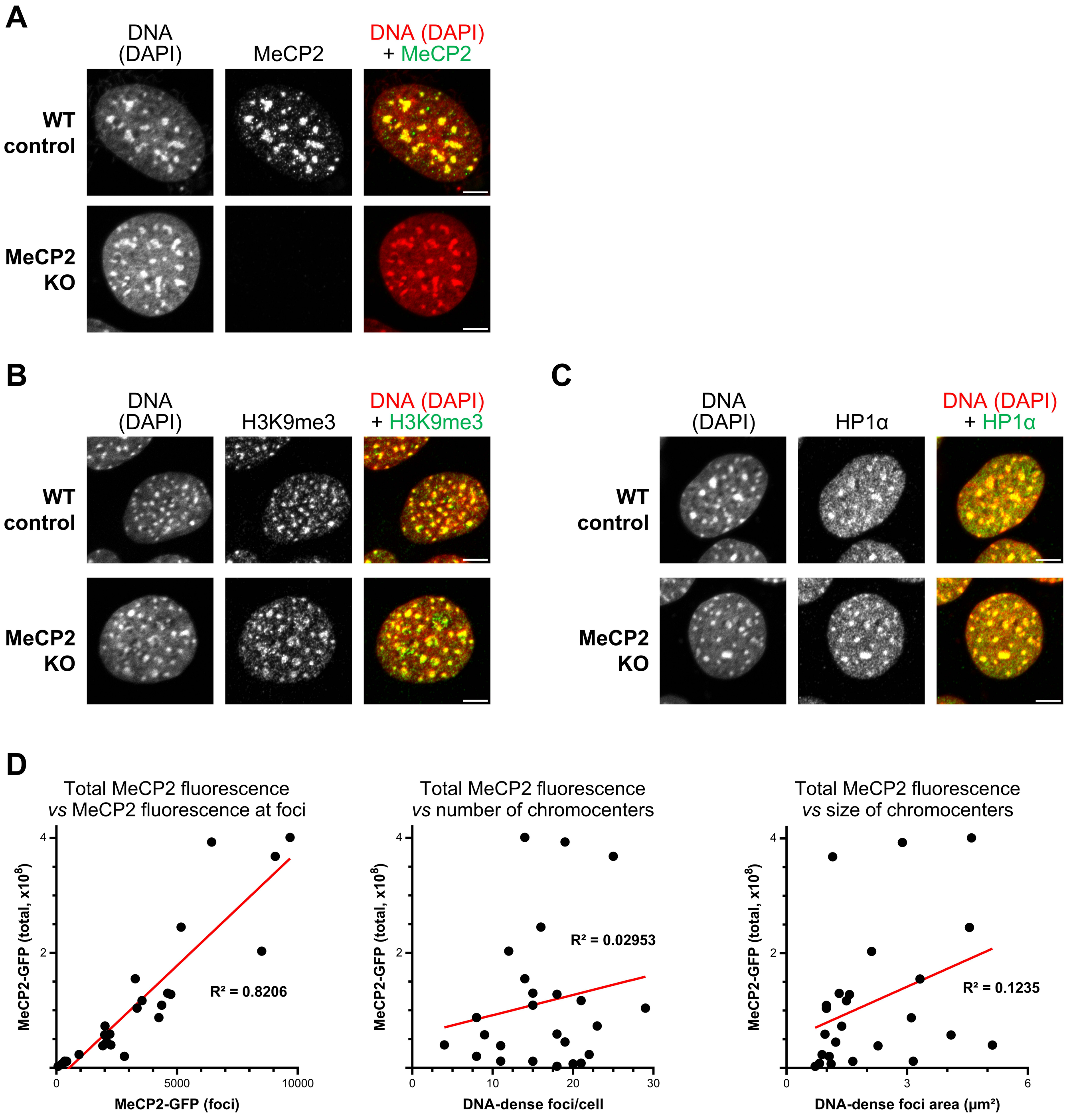

### Figure S6 (related to Figure 3)

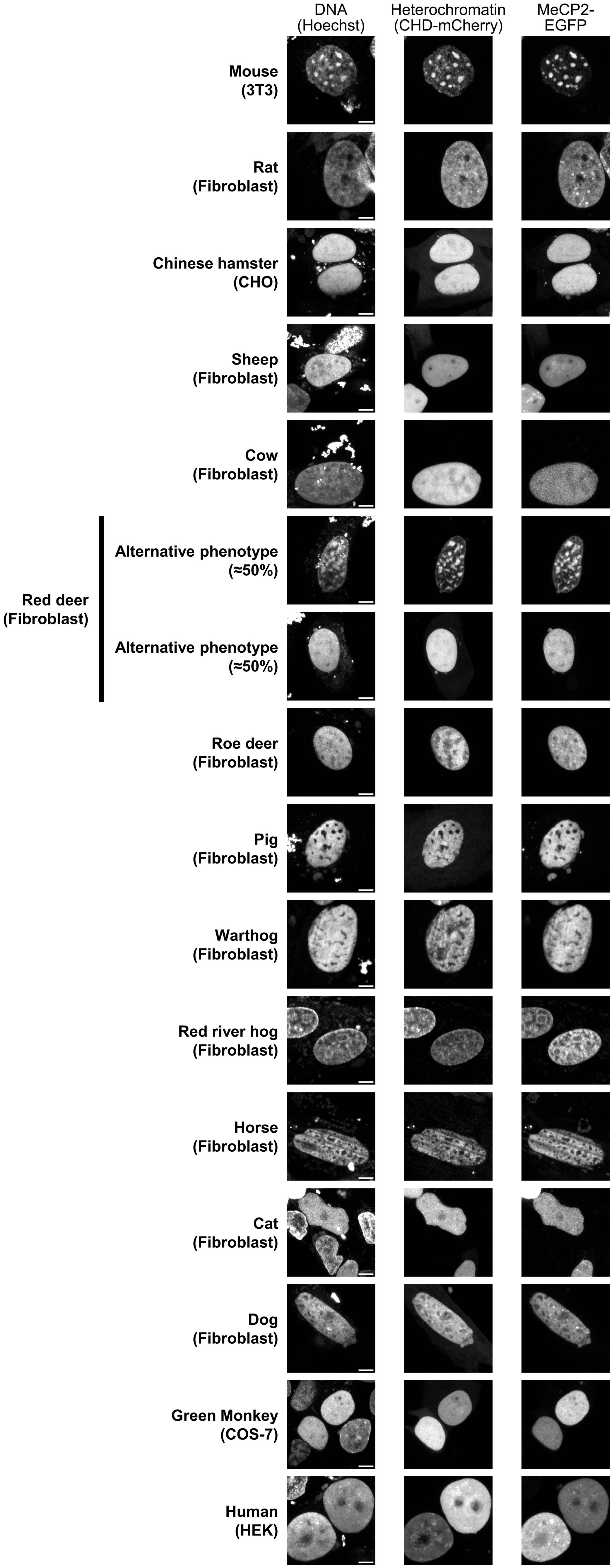

### Figure S7 (related to Figure 3)

```
                     1        10        20        30        40        50        60        70        80
MOUSE                MVAGMLGLREEKSEDQDLQGLRDKPLKFKKAKKDKKEDKEGKHEPLQPSAHHSAEPAEAGKAETSESSGSAPAVPEASAS
RAT                  MVAGMLGLRKEKSEDQDLQGLKEKPLKFKKVKKDKKEDKEGKHEPLQPSAHHSAEPAEAGKAETSESSGSAPAVPEASAS
CHINESE_HAMSTER      MVAGMLGLREEKSEDQDLQGLKDKPLKFKKVKKDKKEDKEGKHEPLQPSAHHSAEPAEAGKAETSESSGSAPAVPEASAS
SHEEP                MVAGMLGLREEKSEEQDLQGLKDKPLKFKKVKKDKKEDKEGKHEPLQPAAHHSAEPAEAGKAETSEGSGSAPAVPEASAS
COW                  MVAGMLGLREEKSEEQDLQGLKDKPLKFKKVKKDKKEDKEGKHEPLQPAAHHSAEPAEAGKAETSEGSGSAPAVPEASAS
RED_DEER             MVAGMLGLREEKSEEQDLQGLKDKPLKFKKVKKDKKEDKEGKHEPLQPAAHHSAEPAEAGKAETSEGSGSAPAVPEASAS
PIG                  MVAGMLGLREEKSEEQDLQGLKDKPLKFKKVKKDKKDDREGKHEPLQPPAHHSAEPAEAGKAETSEGSGSAPAVPEASAS
HORSE                MVAGMLGLREEKSEDQDLQGLKDKPLKFKKVKKDKKEDKEGKHEPLQPPAHHSAEPAEAGKAETSEGSGSAPAVPEASAS
CAT                  MVAGMLGLREEKSEDQDLQGLKDKPLKFKKVKKDKKEDKEGKHEPLQPPAHHSAEPAEAGKAETSEGSGTAPAVPEASAS
DOG                  MVAGMLGLREEKSEDQDLQGLKDKPLKFKKVKKEKKEDKEGKHEPLQPPAHHSAEPAEAGKAETSEGSGSAPAVPEASAS
AFRICAN_GREEN_MONKEY MVAGMLGLREEKSEDQDLQGLKDKPLKFKKVKKDKKEDKEGKHEPVQPSAHHSAEPAEAGKAETSEGSGSAPAVPEASAS
HUMAN                MVAGMLGLREEKSEDQDLQGLKDKPLKFKKVKKDKKEEKEGKHEPVQPSAHHSAEPAEAGKAETSEGSGSAPAVPEASAS

                              90       100       110       120       130       140       150       160
MOUSE                PKQRRSIIRDRGPMYDDPTLPEGWTRKLKQRKSGRSAGKYDVYLINPQGKAFRSKVELIAYFEKVGDTSLDPNDFDFTVT
RAT                  PKQRRSIIRDRGPMYDDPTLPEGWTRKLKQRKSGRSAGKYDVYLINPQGKAFRSKVELIAYFEKVGDTSLDPNDFDFTVT
CHINESE_HAMSTER      PKQRRSIIRDRGPMYDDPTLPEGWTRKLKQRKSGRSAGKYDVYLINPQGKAFRSKVELIAYFEKVGDTSLDPNDFDFTVT
SHEEP                PKQRRSIIRDRGPMYDDPTLPEGWTRKLKQRKSGRSAGKYDVYLINPQGKAFRSKVELIAYFEKVGDTSLDPNDFDFTVT
COW                  PKQRRSIIRDRGPMYDDPTLPEGWTRKLKQRKSGRSAGKYDVYLINPQGKAFRSKVELIAYFEKVGDTSLDPNDFDFTVT
RED_DEER             PKQRRSIIRDRGPMYDDPTLPEGWTRKLKQRKSGRSAGKYDVYLINPQGKAFRSKVELIAYFEKVGDTSLDPNDFDFTVT
PIG                  PKQRRSIIRDRGPMYDDPTLPEGWTRKLKQRKSGRSAGKYDVYLINPQGKAFRSKVELIAYFEKVGDTSLDPNDFDFTVT
HORSE                PKQRRSIIRDRGPMYDDPTLPEGWTRKLKQRKSGRSAGKYDVYLINPQGKAFRSKVELIAYFEKVGDTSLDPNDFDFTVT
CAT                  PKQRRSIIRDRGPMYDDPTLPEGWTRKLKQRKSGRSAGKYDVYLINPQGKAFRSKVELIAYFEKVGDTSLDPNDFDFTVT
DOG                  PKQRRSIIRDRGPMYDDPTLPEGWTRKLKQRKSGRSAGKYDVYLINPQGKAFRSKVELIAYFEKVGDTSLDPNDFDFTVT
AFRICAN_GREEN_MONKEY PKQRRSIIRDRGPMYDDPTLPEGWTRKLKQRKSGRSAGKYDVYLINPQGKAFRSKVELIAYFEKVGDTSLDPNDFDFTVT
HUMAN                PKQRRSIIRDRGPMYDDPTLPEGWTRKLKQRKSGRSAGKYDVYLINPQGKAFRSKVELIAYFEKVGDTSLDPNDFDFTVT

                             170       180       190       200       210       220       230       240
MOUSE                GRGSPSRREQKPPKKPKSPKAPGTGRGRGRPKGSGTGRPKAAASEGVQVKRVLEKSPGKLVVKMPFQASPGGKGEGGGAT
RAT                  GRGSPSRREQKPPKKPKSPKAPGTGRGRGRPKGSGTGRPKAAASEGVQVKRVLEKSPGKLLVKMPFQASPGGKGEGGGAT
CHINESE_HAMSTER      GRGSPSRREQKPPKKPKSPKAPGTGRGRGRPKGSGTGRPKAASSEGVQVKRVLEKSPGKLLVKMPFQASPGGKNEGGGAT
SHEEP                GRGSPSRREQKPPKKPKSPKAPGTGRGRGRPKGSGTTRPKAAASEGVQVKRVLEKSPGKLLVKMPFQAAPGSKAEGGGAT
COW                  GRGSPSRREQKPPKKPKSPKAPGTGRGRGRPKGSGTTRPKAAASEGVQVKRVLEKSPGKLLVKMPFQAAPGSKAEGGGAT
RED_DEER             GRGSPSRREQKPPKKPKSPKAPGTGRGRGRPKGSGTTRPKAAASEGVQVKRVLEKSPGKLLVKMPFQAAPGSKAEGGGAT
PIG                  GRGSPSRREQKPPKKPKSPKAPGTGRGRGRPKGSGTTRPKAAASEGVQVKRVLEKSPGKLLVKMPFQASPGSKAEGGGAT
HORSE                GRGSPSRREQKPPKKPKSPKAPGTGRGRGRPKGSGTARPKAAASEGVQVKRVLEKSPGKLLVKMPFQASPGSKAEGGGAT
CAT                  GRGSPSRREQKPPKKPKSPKAPGTGRGRGRPKGSGTARPKAAASEGVQVKRVLEKSPGKLLVKMPFQASPGSKAEGGGAT
DOG                  GRGSPSRREQKPPKKPKSPKAPGTGRGRGRPKGSGTARPKAATSEGVQVKRVLEKSPGKLLVKMPFQASPGSKAEGGGAT
AFRICAN_GREEN_MONKEY GRGSPSRREQKPPKKPKSPKAPGTGRGRGRPKGSGTTRPKAATSEGVQVKRVLEKSPGKLLVKMPFQTSPGGKAEGGGAT
HUMAN                GRGSPSRREQKPPKKPKSPKAPGTGRGRGRPKGSGTTRPKAATSEGVQVKRVLEKSPGKLLVKMPFQTSPGGKAEGGGAT

                             250       260       270       280       290       300       310       320
MOUSE                TSAQVMVIKRPGRKRKAEADPQAIPKKRGRKPGSVVAAAAAEAKKKAVKESSIRSVHETVLPIKKRKTRETVSIEVKEVV
RAT                  TSAQVMVIKRPGRKRKAEADPQAIPKKRGRKPGSVVAAAAAEAKKKAVKESSIRSVQETVLPIKKRKTRETVSIEVKEVV
CHINESE_HAMSTER      TSAQVMVIKRPGRKRKAEADPQAIPKKRGRKPGSVVAAAAAEAKKKAVKESSIRSVHETVLPIKKRKTRETVSIEVKEVV
SHEEP                TSAQVMVIKRPGRKRKAEADPQAIPKKRGRKPGSVVAAAAAEAKKKAVKESSIRSVQETVLPIKKRKTRETVSIEVKEVV
COW                  TSAQVMVIKRPGRKRKAEADPQAIPKKRGRKPGSVVAAATAEAKKKAVKESSIRSVQETVLPIKKRKTRETVSIEVKEVV
RED_DEER             TSAQVMVIKRPGRKRKAEADPQAIPKKRGRKPGSVVAAAAAEAKKKAVKESSIRSVQETVLPIKKRKTRETVSIEVKEVV
PIG                  TSAQVMVIKRPGRKRKAEADPQAIPKKRGRKPGSVVAAAAAEAKKKAVKESSIRSVQETVLPIKKRKTRETVSIEVKEVV
HORSE                TSAQVMVIKRPGRKRKAEADPQAIPKKRGRKPGSVVAAAAAEAKKKAVKESSIRSVQETVLPIKKRKTRETVSIEVKEVV
CAT                  TSAQVMVIKRPGRKRKAEADPQAIPKKRGRKPGSVVAAAAAEAKKKAVKESSIRSVQETVLPIKKRKTRETVSIEVKEVV
DOG                  TSAQVMVIKRPGRKRKAEADPQAIPKKRGRKPGSVVAAAAAEAKKKAVKESSIRSVQETVLPIKKRKTRETVSIEVKEVV
AFRICAN_GREEN_MONKEY TSTQVMVIKRPGRKRKAEADPQAIPKKRGRKPGSVVAAAAAEAKKKAVKESSIRSVQETVLPIKKRKTRETVSIEVKEVV
HUMAN                TSTQVMVIKRPGRKRKAEADPQAIPKKRGRKPGSVVAAAAAEAKKKAVKESSIRSVQETVLPIKKRKTRETVSIEVKEVV

                             330       340       350       360       370       380       390
MOUSE                KPLLVSTLGEKSGKGLKTCKSPGRKSKESSPKGRSSSAS..SPPKKEHHHHHHSESTKAPMPLLPSPP..PPEPESSED
RAT                  KPLLVSTLGEKSGKGLKTCKSPGRKSKESSPKGRSSSAS..SPPKKEHHHHHHAESPKAPMPLLPPPP..PPEPQSSED
CHINESE_HAMSTER      KPLLVSTLGEKSGKGLKTCKSPGRKSKESSPKGRSSSSASSPPKKEHHHHHHSESPKAPMPLLPPPPPPPPEPESSED
SHEEP                KPLLVSTLGEKSGKGLKTCKSPGRKSKESSPKGRSGSAS..SPPKKEHHHHHHVEPPKAPAPLLLPPPPPPPEPQSSED
COW                  KPLLVSTLGEKSGKGLKTCKSPGRKSKESSPKGRSGSAS..SPPKKEHHHHHHVEPPKAPAPLLLPPPPPPPEPQSSED
RED_DEER             KPLLVSTLGEKSGKGLKTCKSPGRKSKESSPKGRSGSAS..SPPKKEHHHHHHAEPPKAPAPLLLPPPPPPPEPQSSED
PIG                  KPLLVSTLGEKSGKGLKTCKSPGRKSKEGSA.......S..SPPKKEHHHHHHAEPPKAPAPLLPPPPPPPPEPQSSED
HORSE                KPLLVSTLGEKSGKGLKTCKSPGRKSKESSPKGRSSTS..SPPKKEHHHHHHAEPPRAPAPLLPPPPPPPPEPQSSED
CAT                  KPLLVSTLGEKSGKGLKTCKSPGRKSKESSPKGRGGSAS..SPPKKEHHHHHHSEPPKAPAPLLPPPPPPPPEPQSSED
DOG                  KPLLVSTLGEKSGKGLKTCKSPGRKSKESSPKGRSSSAS..SPPKKEHHHHHHSEPPKAPAPLLPPPPPPPPEPQSSED
AFRICAN_GREEN_MONKEY KPLLVSTLGEKSGKGLKTCKSPGRKSKESSPKGRSSSAS..SPPKKEHHHHHHSESPKAPVPLLPPLPPPPPEPESSED
HUMAN                KPLLVSTLGEKSGKGLKTCKSPGRKSKESSPKGRSSSAS..SPPKKEHHHHHHSESPKAPVPLLPPLPPPPPEPESSED

                     400       410       420       430       440                450       460
MOUSE                PISPPEPQDLSSSICKEEKMPRGGSLESDGCPKEPAKTQPMVAT.........TTTVAEKYKHRGEGERKDIVSSSMPRPN
RAT                  PISPPEPQDLSSSICKEEKMPRAGSLESDGCPKEPAKTQPMVAAAATTTTTTTTTVAEKYKHRGEGERKDIVSSSMPRPN
CHINESE_HAMSTER      PISPPEPQDLSSSICKEEKMPRGGSLESDGCPKEPAKTQPMVAAT........TATVAEKYKHRGEGERKDIVSSSMPRPN
SHEEP                PASPPEPQDLSSSVCKEEKMPRAGSLESDGCPKEPAKTQPALATA.........APATEKYKHRGEGERKDIVSSSMPRPN
COW                  PASPPEPQDLSSSVCKEEKMPRAGSLESDGCPKEPAKTQPALATA.........APATEKYKHRGEGERKDIVSSSMPRPN
RED_DEER             PASPPEPQDLSSSICKEEKMPRAGSLESDGCPKEPAKTQPTLATA.........APATEKYKHRGEGERKDIVSSSMPRPN
PIG                  PASPPEPQDLSSNVCREEKMPRAGSLESDGCPKEPAKTQPAVATA.........ATAAEKYKHRGEGERKDIVSSSMPRPN
HORSE                PTSPPEPQDLSSSVCKEEKMPRGGSLESDGCPKEPAKTQPAVATA.........ATAAEKYKHRGEGERKDIVSSSMPRPN
CAT                  PASPPEPQDLSSGVCKEEKMPRGGSLESDGCPKEPAKTQPAVATA.........ATAAEKYKHRGEGERKDIVSSSMPRPN
DOG                  PASPPEPQDLSSGVCKEEKMARGGSLESDGCPKEPAKTQPTVATA.........ATAADKYKHRGEGERKDIVSSSMPRPN
AFRICAN_GREEN_MONKEY PTSPPEPQDLSSSVCKEEKMPRGGSLESDGCPKEPAKTQPAVATA.........ATAAEKYKHRGEGERKDIVSSSMPRPN
HUMAN                PTSPPEPQDLSSSVCKEEKMPRGGSLESDGCPKEPAKTQPAVATA.........ATAAEKYKHRGEGERKDIVSSSMPRPN

                    470       480
MOUSE                REEPVDSRTPVTERVS
RAT                  REEPVDSRTPVTERVS
CHINESE_HAMSTER      REEPVDSRTPVTERVS
SHEEP                REEPVDSRTPVTERVS
COW                  REEPVDSRTPVTERVS
RED_DEER             REEPVDSRTPVTERVS
PIG                  REEPVDSRTPVTERVS
HORSE                REEPVDSRTPVTERVS
CAT                  REEPVDSRTPVTERVS
DOG                  REEPVDSRTPVTERVS
AFRICAN_GREEN_MONKEY REEPVDSRTPVTERVS
HUMAN                REEPVDSRTPVTERVS
```

### Figure S8 (related to Figure 3)

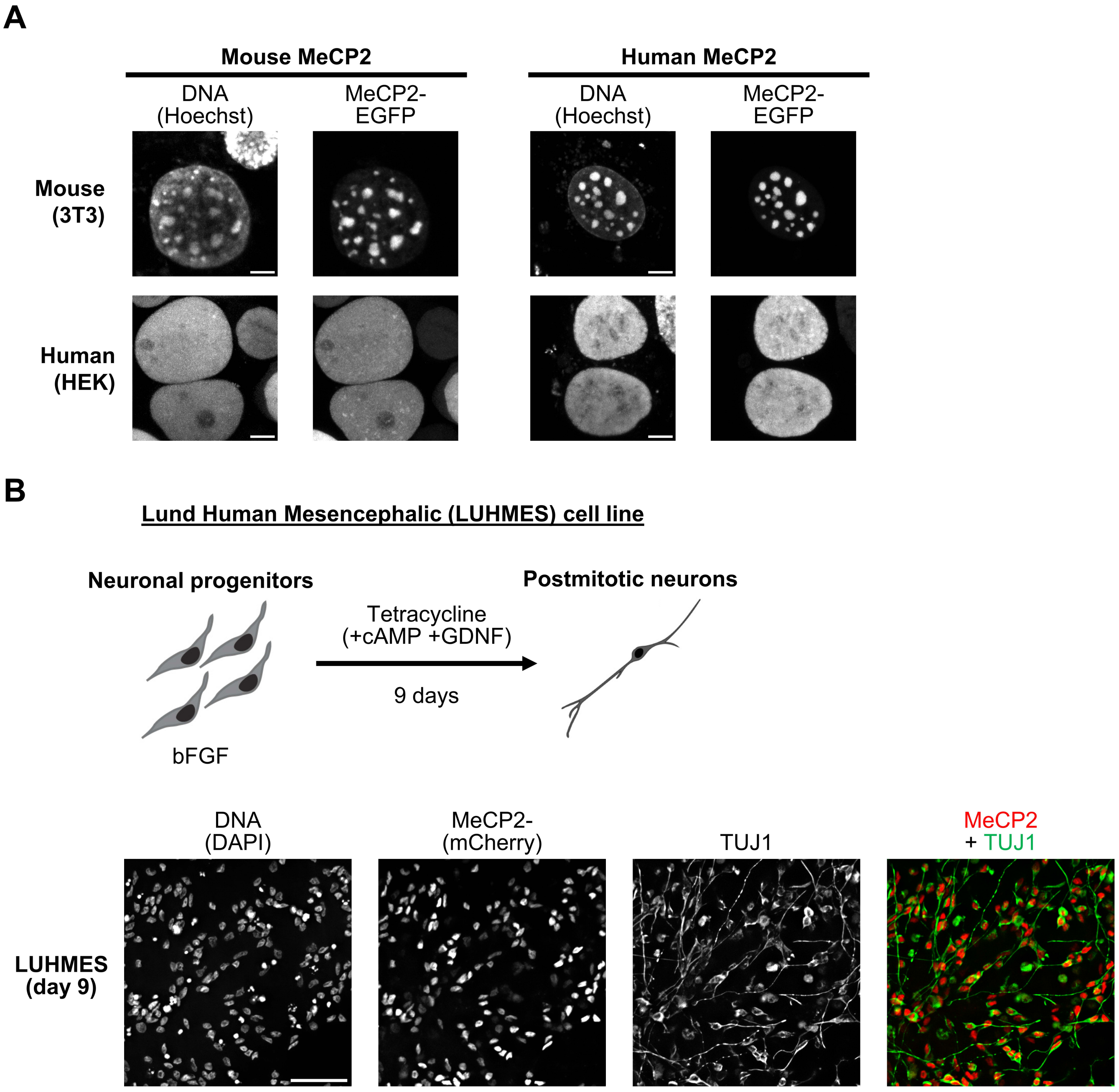

### Figure S9 (related to Figure 4)

**A**

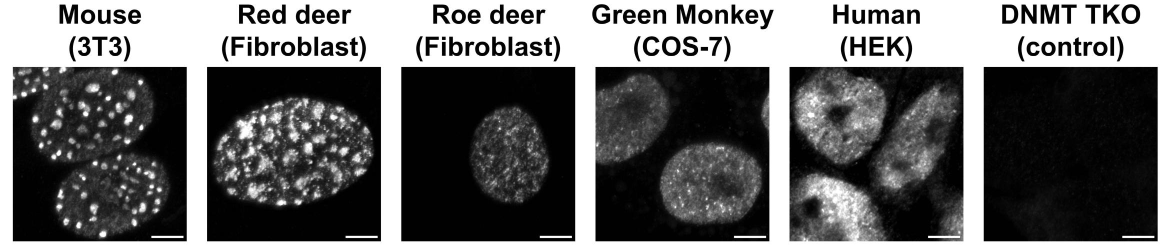


**B**

5’ CAAGACGAAAGGATGTCTGAATCCCCTGTGGAGACCACAGAGAAAGACCTAGTTCCCCACCTCATC
3’ GTTCTGCTTTCCTACAGACTTAGGGGACACCTCTGGTGTCTCTTTCTGGATCAAGGGGTGGAGTAG

GCGACCGGAGGCCTCACATCCTTTGAAAACTCCAGAGGTACGCGGAGATCAGTGCCTCCAAAGGAG
CGCTGGCCTCCGGAGTGTAGGAAACTTTTGAGGTCTCCATGCGCCTCTAGTCACGGAGGTTTCCTC

ACGATGCCTGACTCCTCGTGAAACTTGATAGGAGTCCCAGGATTCCTGTGGCACGTGGAAAGGGAC
TGCTACGGACTGAGGAGCACTTTGAACTATCCTCAGGGTCCTAAGGACACCGTGCACCTTTCCCTG

CCTTGGTCTCCCGCCTCAGCTGGAGAGGCGTCCCAATTGCCCTGCCAAGCCTCGAGGAGAATCCCG
GGAACCAGAGGGCGGAGTCGACCTCTCCGCAGGGTTAACGGGACGGTTCGGAGCTCCTCTTAGGGC

AGTTGTCCCTCGCAACTAGGCAGGAGTCCTGACGTCGCTGAAGAAACACGTGTGTGGAAGGGCCAT
TCAACAGGGAGCGTTGATCCGTCCTCAGGACTGCAGCGACTTCTTTGTGCACACACCTTCCCGGTA

CCCCGTCGTAACTCGAGAATATACCCCAGGTTCCCGCCGCAACTCGAGAAAAACCATGAGACTTCC
GGGGCAGCATTGAGCTCTTATATGGGGTCCAAGGGCGGCGTTGAGCTCTTTTTGGTACTCTGAAGG

CCCTCGCCGCGAGATGAGGCCCGATTCCCCTGCACTGCGTGCAGAGCAATTCCGTGTTGCACATCA
GGGAGCGGCGCTCTACTCCGGGCTAAGGGGACGTGACGCACGTCTCGTTAAGGCACAACGTGTAGT

CACATGAAAGGAGCCTTGATTTCCTTGATGGCACTCCAGAGAAACCCCAAGAACACTGTTTCAAGG
GTGTACTTTCCTCGGAACTAAAGGAACTACCGTGAGGTCTCTTTGGGGTTCTTGTGACAAAGTTCC

CTAGAGGGATCCTGAGGTCACTGTAGCAACACGAAAGAGCTCCGTGGACCAAAAATCAACTCGAGA
GATCTCCCTAGGACTCCAGTGACATCGTTGTGCTTTCTCGAGGCACCTGGTTTTTAGTTGAGCTCT

TGAGAGGTTAGTCCCTGGCTTCGACTCCAGAGGAATACCACCTTACCACAAGCACCTCAAGAGGAG
ACTCTCCAATCAGGGACCGAAGCTGAGGTCTCCTTATGGTGGAATGGTGTTCGTGGAGTTCTCCTC

GCTTCTCTCAGCTCTAGGTATGTGAGAGGGACCCTGAGTTTGCGGCCTCAAGTGGAATGGACACC 3’
CGAAGAGAGTCGAGATCCATACACTCTCCCTGGGACTCAAACGCCGGAGTTCACCTTACCTGTGG 5’

**C**

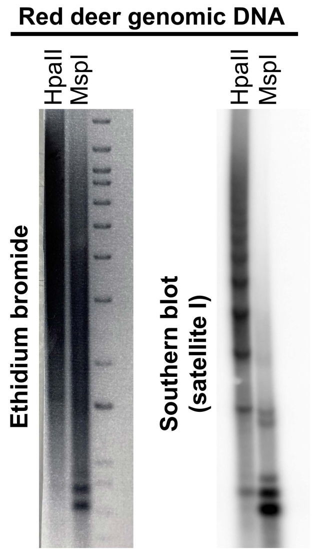


**D**

5’ GGACCTGGAATATGGCGAGAAAACTGAAAATCACGGAAAATGAGAAATACACACTTTAGG
3’ CCTGGACCTTATACCGCTCTTTTGACTTTTAGTGCCTTTTACTCTTTATGTGTGAAATCC

ACGTGAAATATGGCGAGGAAAACTGAAAAAGGTGGAAAATTTAGAAATGTCCACTGTAGG
TGCACTTTATACCGCTCCTTTTGACTTTTTCCACCTTTTAAATCTTTACAGGTGACATCC

ACGTGGAATATGGCAAGAAAACTGAAAATCATGGAAAATGAGAAACATCCACTTGACGAC
TGCACCTTATACCGTTCTTTTGACTTTTAGTACCTTTTACTCTTTGTAGGTGAACTGCTG

TTGAAAAATGACGAAATCACTAAAAAACGTGAAAAATGAGAAATGCACACTGAA 3’
AACTTTTTACTGCTTTAGTGATTTTTTGCACTTTTTACTCTTTACGTGTGACTT 5’
